# DNA Fountain enables a robust and efficient storage architecture

**DOI:** 10.1101/074237

**Authors:** Yaniv Erlich, Dina Zielinski

## Abstract

DNA is an attractive medium to store digital information. Here, we report a storage strategy, called DNA Fountain, that is highly robust and approaches the information capacity per nucleotide. Using our approach, we stored a full computer operating system, movie, and other files with a total of 2.14 × 10^6^ bytes in DNA oligos and perfectly retrieved the information from a sequencing coverage equivalent of a single tile of Illumina sequencing. We also tested a process that can allow 2.18 × 10^15^ retrievals using the original DNA sample and were able to perfectly decode the data. Finally, we explored the limit of our architecture in terms of bytes per molecules and obtained a perfect retrieval from a density of 215Petabyte/gram of DNA, orders of magnitudes higher than previous techniques.

## Text

Humanity is currently producing data at exponential rates, creating a demand for better storage devices. DNA is an excellent medium for data storage with demonstrated information density of petabytes of data per gram, high durability, and evolutionarily optimized machinery to faithfully replicate this information ^1,2^. Recently, a series of proof-of-principle experiments have demonstrated the value of DNA as a storage medium ^3–9^.

To better understand its potential, we explored the Shannon information capacity ^10,11^ of DNA storage ^12^. This measure sets a tight upper bound on the amount of information that can be reliably stored in each nucleotide. In an ideal world, the information capacity of each nucleotide could reach 2bits, since there are four possible options. However, DNA encoding faces several practical limitations. First, not all DNA sequences are created equal ^13,14^. Biochemical constraints dictate that DNA sequences with high GC content or long homopolymer runs (e.g. AAAAAA…) should be avoided as they are difficult to synthesize and prone to sequencing errors. Second, oligo synthesis, PCR amplification, and decay of DNA during storage can induce uneven representation of the oligos ^7,15^. This might result in dropout of a small fraction of oligos that will not be available for decoding. In addition to biochemical constraints, oligos are sequenced in a pool and necessitate indexing to infer their order, which further limits the number of available nucleotides for encoding information. Quantitative analysis shows that the biochemical constraints reduce the coding potential of each nucleotide to 1.98bits. After combining the expected dropout rates and barcoding demand, the overall Shannon information capacity of a DNA storage device is around 1.83bits per nucleotide for a range of practical architectures ^12^ (**Supplementary Figure 1-5; Supplementary Table. 1-2**).

Previous studies of DNA storage realized about half of the Shannon information capacity of DNA molecules and in some cases reported challenges to perfectly retrieve the information (**Table 1**). For example, two previous studies attempted to address oligo dropout by dividing the original file into overlapping segments so that each input bit is represented by multiple oligos ^4,6^. However, this repetitive coding procedure generates a loss of information content and is poorly scalable and (**Supplementary Figure 6**). In both cases, these studies reported small gaps in the retrieved information ^3,4,6^. Other studies explored the usage of Reed-Solomon (RS) code on small blocks of the input data in order to recover dropouts ^5,7^. While these studies were able to perfectly retrieve the data, they were still far from realizing the capacity. Moreover, testing this strategy on a large file size highlighted difficulties in decoding the data due to local correlations and large variations in the dropout rates within each protected block ^7^, which is a known issue of blocked RS codes ^16,17^. Only after employing a complex multi-step procedure and high sequencing coverage, the study was able to rescue a sufficient number of oligos. Taken together, these results inspired us to seek a coding strategy that can better utilize the information capacity of DNA storage devices while showing higher data retrieval reliability.

**Table 1:** Comparison of DNA storage coding schemes and experimental results. For consistency, describes only schemes that were empirically tested with pooled oligo synthesis and high throughput sequencing data. The schemes are presented chronologically on the basis of date of publication ^(1)^Coding potential is the maximal information content of each nucleotide before indexing or error correcting ^(2)^Redundancy denotes the excess of synthesized oligos to provide robustness to dropouts ^(3)^The proposed strategy to recover oligo dropouts (RS, Reed-Solomon codes) ^(4)^The presence of error correcting/detection code to handle synthesis and sequencing errors ^(5)^Whether all information was recovered without any error ^(6)^The input information in bits divided by the number of overall DNA bases requested for sequencing (excluding adapter annealing sites) ^(7)^The ratio between the net information density and the Shannon Capacity of the channel ^(8)^Physical density is the actual ratio of the number of bytes encoded and minimal weight of the DNA library used to retrieve the information. This information was not available for studies by Bornholt et al. and Blawat et al. P: petabyte. See **Supplementary Material** for more information.

|  | Church et al.<sup>3</sup> | Goldman et al.<sup>4</sup> | Grass et al.<sup>5</sup> | Bornholt et al.<sup>6</sup> | Blawat et al.<sup>7</sup> | This work |
| --- | --- | --- | --- | --- | --- | --- |
| Input data[Mbyte] | 0.65 | 0.75 | 0.08 | 0.15 | 22 | 2.15 |
| Coding potential<sup>(1)</sup> [bits/nt] | 1 | 1.58 | 1.78 | 1.58 | 1.6 | 1.98 |
| Redundancy<sup>(2)</sup> | 1 | 4 | 1 | 1.5 | 1.13 | 1.07 |
| Robustness to dropouts<sup>(3)</sup> | No | Repetition | RS | Repetition | RS | Fountain |
| Error correction/detection<sup>(4)</sup> | No | Yes | Yes | No | Yes | Yes |
| Full recovery<sup>(5)</sup> | No | No | Yes | No | Yes | Yes |
| Net information density<sup>(6)</sup> [bits/nt] | 0.83 | 0.33 | 1.14 | 0.88 | 0.92 | 1.57 |
| Realized capacity<sup>(7)</sup> | 45% | 18% | 62% | 48% | 50% | 86% |
| Physical density<sup>(8)</sup> [PB/g] | 1.28 | 2.25 | 0.005 | - | - | 214 |

We devised a strategy for DNA storage devices, called DNA Fountain, that approaches the Shannon capacity while providing strong robustness against data corruption. Our strategy harnesses fountain codes ^18,19^, which allows reliable and effective unicasting of information over channels that are subject to dropouts, such as mobile TV ^20^. In our design, we carefully adapted the power of fountain codes to overcome both oligo dropouts and the biochemical constraints of DNA storage. Our encoder works in three steps (**Fig. 1**) ^12^: First, it preprocesses a binary file into a series of non-overlapping segments of a certain length. Next, it iterates over two *computational* steps: Luby Transform and screening. The Luby Transform sets the basis for fountain codes. Basically, it packages data into any desired number of short messages, called droplets, by selecting a random subset of segments from the file using a special distribution (**Supplementary Figure 7**) and adding them bitwise together under a binary field. The droplet contains two pieces of information: a data payload part that holds the result of the addition procedure and a short, fixed-length seed. This seed corresponds to the state of the random number generator of the transform during the droplet creation and allows the decoder algorithm to infer the identities of the segments in the droplet. Theoretically, reversing the Luby Transform to recover the file is possible using a highly efficient algorithm by collecting any subset of droplets as long as the accumulated size of droplets is slightly bigger than the size of the original file. We apply one round of the transform in each iteration to create a single droplet. Next, the algorithm moves to the droplet screening stage. This stage is not part of the original fountain code design. It completely realizes the coding potential of each nucleotide under the biochemical constraints to reach 1.98bit/nt. In screening, the algorithm translates the binary droplet to a DNA sequence by converting {00,01,10,11} to {A,C,G,T}, respectively. Then, it screens the sequence for the desired biochemical properties of GC content and homopolymer runs. If the sequence passes the screen, it is considered valid and added to the oligo design file; otherwise, the algorithm simply trashes the droplet. Since the Luby Transform can create any desired number of droplets, we keep iterating over the droplet creation and screening steps until a sufficient number of valid oligos are generated. In practice, we recommend 5%-10% more oligos than input segments ^12^. Searching for valid oligos scales well with the size of the input file and is economical for various oligo lengths within and beyond current synthesis limits ^12^ (**Table S3**).

**Figure 1:**
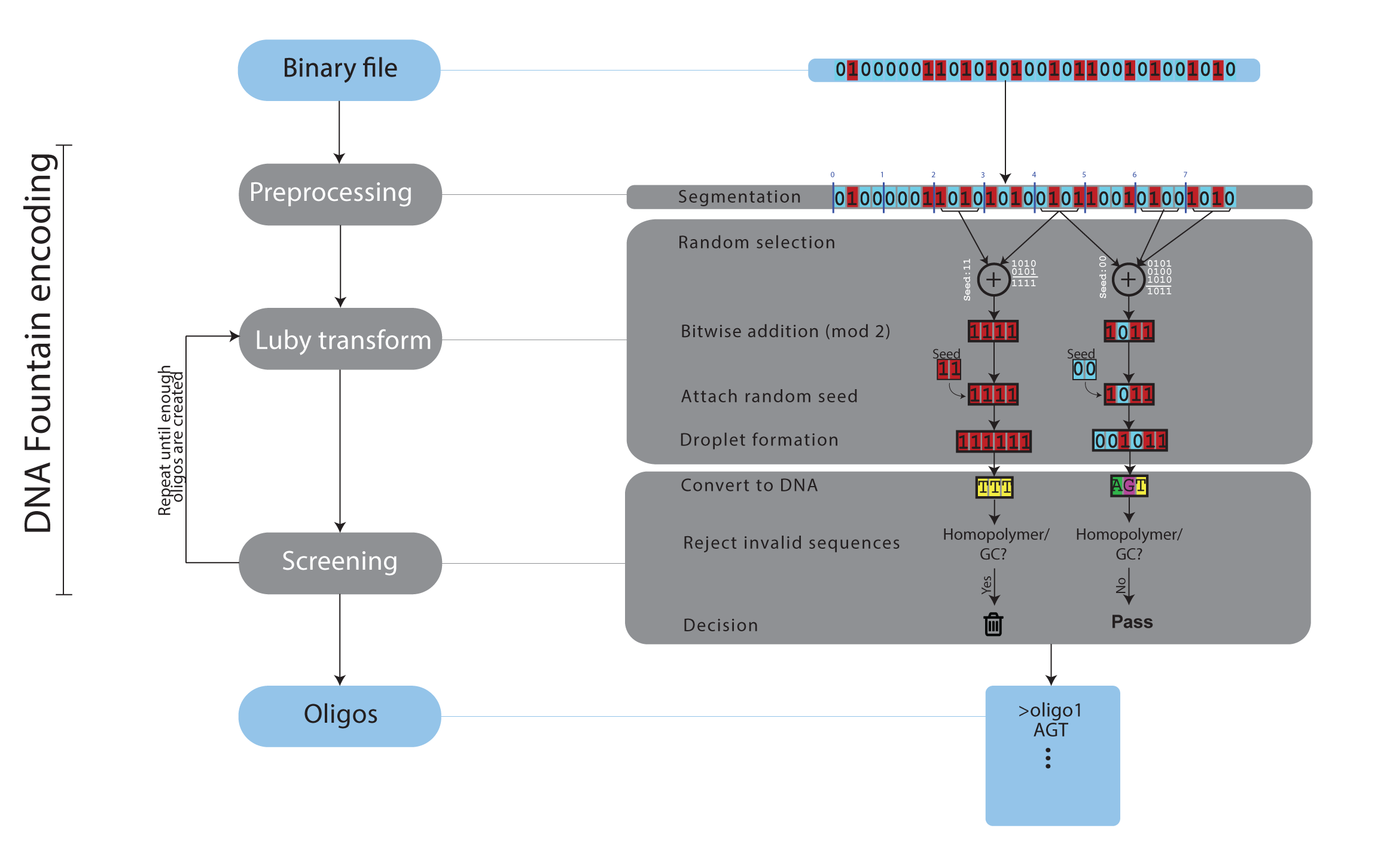
DNA fountain encoding. Left: three main algorithmic steps. Right: an example with a small file of 32bits. For simplicity, we partitioned the file into 8 segments of 4bits. The seeds are represented as 2bit numbers and are presented for display purposes only. See **Supplementary Material** for the full details of each algorithmic step.

We used DNA Fountain to encode a single compressed file of 2,146,816 bytes in a DNA oligo pool. The input data was a tarball that packaged several files, including a complete graphical operating system of 1.4Mbyte and a $50 Amazon gift card ^12^ (**Figs. 2A; S8**). We split the input tarball into 67,088 segments of 32bytes and iterated over the steps of DNA Fountain to create valid oligos. Each droplet was 38bytes (304bits): 4bytes of the random number generator seed, 32bytes for the data payload, and 2bytes for a Reed-Solomon error correcting code, to reject erroneous oligos in low coverage conditions. With this seed length, our strategy supports encoding files of up to 500Mbyte ^12^. The DNA oligos had a length of (304/2)=152nt and were screened for homopolymer runs of ≤3nt and GC content of 45%-55%. We instructed DNA Fountain to generate 72,000 oligos, yielding a redundancy of (72,000/67,088-1)=7%. We selected this number of oligos due to the price structure offered by the manufacturer, allowing us to maximize the number of oligos per dollar. Finally, we added upstream and downstream annealing sites for Illumina adapters, making our final oligos 200nt long (**Figs. 2B; S9**). Encoding took 2.5min on a single CPU of a standard laptop. Importantly, we achieved an information density of 1.57bit/nt, only 14% from the Shannon capacity of DNA storage and 60% more than previous studies with a similar scale of data (**Table 1**).

**Figure 2:**
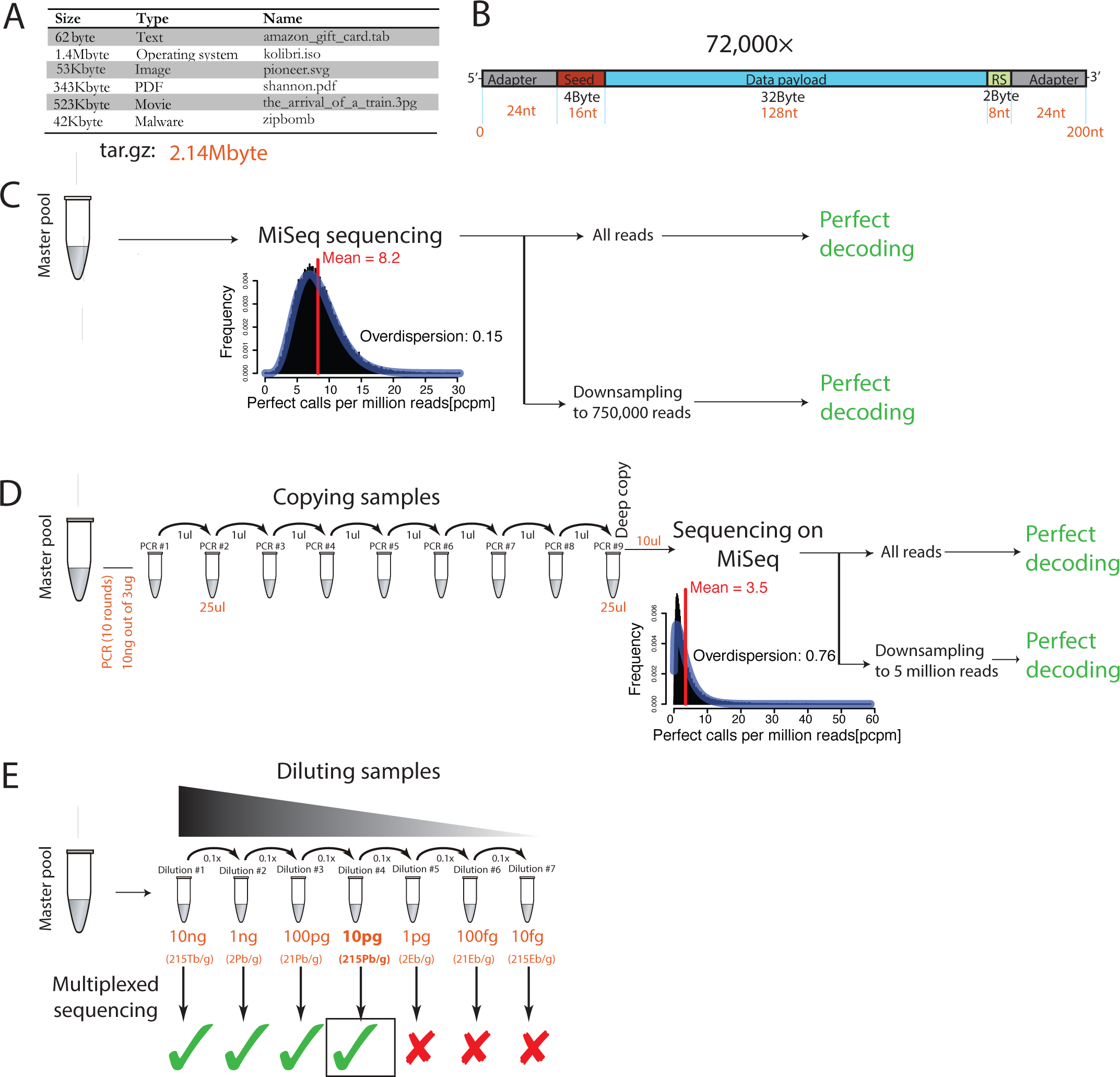
Experimental setting and results for storing data on DNA. **(A)** The input files for encoding, size, and type. The total amount of data was 2.14Mbyte after compression **(B)** The structure of the oligos. Black labels: length in bytes. Red: length in nucleotides. RS: Reed-Solomon error correcting code **(C)** experimental results of the master pool **(C+D)** The histogram displays the frequency of perfect calls per million (pcpm) sequenced reads. Red: mean. Blue: negative binomial fit to the pcpm **(D)** Experimental procedures of deep copying of the oligo pool **(E)** Serial dilution experiment. The weight corresponds to the input DNA library that underwent PCR and sequencing. Green: correctly decoded. Red: decoding failure.

Sequencing and decoding the oligo pool fully recovered the entire input file with zero errors (**Fig. 2C**). To retrieve the information, we PCR-amplified the oligo pool and sequenced the DNA library on one MiSeq flowcell with 150 paired-end cycles, which yielded 32 million reads. We employed a pre-processing strategy that prioritizes reads that are more likely to represent high quality oligos ^12^. Since not all oligos are required for the decoding due to redundancy, this procedure reduces the exposure to erroneous oligos. Then, the decoder scanned the reads, recovered the binary droplets, rejected droplets with errors based on the Reed-Solomon code, and employed a message passing algorithm to reverse the Luby Transform and obtain the original data ^12^.

In practice, decoding took approximately 9min with a Python script on a single CPU of a standard laptop (**Supplementary Movie 1**). The decoder recovered the information with 100% accuracy after observing only 69,870 oligos out of the 72,000 in our library (**Supplementary Figure 10**). To further test the robustness of our strategy, we down-sampled the raw Illumina data to 750,000 reads, equivalent to one tile of an Illumina MiSeq flow cell. This procedure resulted in 1.3% oligo dropout from the library. Despite these limitations, the decoder was able to perfectly recover the original 2.1Mbyte in twenty out of twenty random down-sampling experiments. These results indicate that beyond its high information density, DNA Fountain also reduces the amount of sequencing required for data retrieval, which is beneficial when storing large-scale information.

DNA Fountain can also perfectly recover the file after creating a deep copy of the sample. One of the caveats of DNA storage is that each retrieval of information consumes an aliquot of the material. Copying the oligo library with PCR is possible, but this procedure introduces noise and induces oligo dropout. To further test the robustness of our strategy, we created a deep copy of the file by propagating the sample through nine serial PCR amplifications (**Fig. 2D**). The first PCR reaction used 10ng of material out of the 3ug master pool. Each subsequent PCR reaction consumed 1ul of the previous PCR reaction and employed 10 cycles in each 25ul reaction. We sequenced the final library using one run on the Illumina MiSeq.

Overall, this recursive PCR reflects one full arm of an exponential process that theoretically could generate 300×25^9^×2 = 2.28 quadrillion copies of the file by repeating the same procedure with each aliquot (**Supplementary Figure 11**). As expected, the quality of the deep copy was substantially worse than the initial experiment with the master pool. The average coverage per oligo dropped from an average of 7.8 perfect calls for each oligo per million reads (pcpm) to 4.5pcpm in the deep copy. In addition, the deep copy showed much higher skewed representation with a negative binomial overdispersion parameter (1/size) of 0.76 compared to 0.15 in the master pool. Despite the lower quality, the DNA Fountain decoder was able to fully recover the file without a single error with the full sequencing data. Aftter down-sampling the sequencing data to five million reads, resulting in an approximate dropout rate of 1. 0%, we were able to perfectly recover the file in ten out of ten trials. These results suggest that with DNA fountain, DNA storage can be copied virtually an unlimited number of times while preserving the data integrity of the sample.

Next, we explored the maximal achievable physical density using DNA Fountain. The pioneering study by Church et al. predicted that DNA storage could theoretically achieve an information density of 680 Pbyte (P: peta-; 10^15^) per gram of DNA, assuming the storage of 100 molecules per oligo sequence ^3^. However, previous DNA storage experiments have never attempted to reach this high density and in practice tested data retrieval from libraries with densities that are lower by a factor of at least 300* (**Table 1**). To test the maximal physical density, we sequentially diluted our library by seven orders of magnitude from 10ng of DNA to 10fg (femotogram; 10^-15^) of DNA (**Fig. 2E**). Under a prefect synthesis process (no synthesis errors and/or fragmented DNA molecules), the first dilution (10ng) corresponds to ~10^6^ copies per oligo and a density of ~2Tbyte/g, whereas the last dilution corresponds to ~1 copy per oligo and ~200Ebyte/g (E: exa-; 10^18^). We PCR amplified all libraries using an identical strategy to keep all conditions uniform and sequenced the libraries using two multiplexed Illumina rapid runs, which yielded similar number of reads and quality metrics ^12^.

We were able to perfectly retrieve the information from a physical density of 215Pb/g. This density is over two orders of magnitude higher than previous reports and close to the theoretical prediction by Church et al. At this density, the input weight of the library was 10pg and each oligo was represented by approximately 1300 molecules on average (**Table S4**). We found about a 4% dropout rate, close to the limit of our decoder. For the lower input weights, the libraries had substantially more oligo dropouts, ranging from 62% for the 1pg condition (~2Ebyte/g) to 87% for the 10fg condition (~200Ebyte/g). A more aggressive error correcting capability than DNA Fountain is unlikely to dramatically improve the physical density. We tested decoding of the low-weight libraries (<10pg) under the unrealistic assumption of a decoder that can correct any number of indels and substitutions as long as a very short stretch (15nt) of the read is still intact ^12^. Even this aggressive error correction failed to bring the dropout rates of the 1pg library below 30%. Therefore, these results suggest that the current design gets close to the maximal physical density permitted by the stochastic bottleneck induced by PCR amplification of highly complex libraries using a small number of DNA molecules.

DNA as a storage medium is gaining increased attention. Microsoft has announced the beginning of large-scale experiments to store about 100Mbyte of data in DNA and the United States Intelligence Advanced Research Projects Activity (IARPA) and the Library of Congress also expressed interest in this domain ^21^. The main issue is still the high cost of DNA storage. However, these costs reflect the relatively low throughput of manufacturing high quality oligos for traditional synthetic biology applications that are sensitive to errors ^22^. This is not the case for DNA storage. In our experiments, DNA Fountain was able to perfectly decode the data from conditions that are well below the initial quality and quantity of the oligo manufacturer while approaching the information capacity. With these results, we envision a cost-effective DNA storage architecture that relies on quick-and-dirty oligo synthesis that consume less machine time and reagents and a coding strategy, such as DNA Fountain, to handle the chemical imperfections using strong information-theoretic tools.

## Acknowledgments

Y.E. holds a Career Award at the Scientific Interface from the Burroughs Wellcome Fund. This study was supported by a generous gift from Andria and Paul Heafy to the Erlich Lab. We thank P. Smibert and M. Stoeckius for technical assistance with oligo design, N. Abe and S. Pescatore for sequencing operations, A. Gordon for creating the movie, and S. Zaaijer and N. Sanjana for useful comments and discussions. Code and sequencing data are available on https://github.com/TeamErlich/dna-fountainunderGPLv3.

## Materials and Methods

### 1.1 The Shannon Information Capacity of DNA Storage

#### 1.1.1 Constraints in encoding DNA information

DNA storage is basically a communication channel. We transmit information over the channel by synthesizing DNA oligos. We receive information by sequencing the oligos and decoding the sequencing data. The channel is noisy due to various experimental factors, including DNA synthesis imperfections, PCR dropout, stutter noise, degradation of DNA molecules over time, and sequencing errors. Different from classical information theoretic channels (e.g. the binary symmetric channel) where the noise is identical and independently distributed, the error pattern in DNA heavily depends on the input sequence. Previous studies of error patterns identified that homopolymer runs and GC content are major determinants of synthesis and sequencing errors[13, 14, 23, 24, 25, 26, 27]. For example, Schwartz et al.[13] reported that oligos with a GC content above 60% exhibit high dropout rates and that most PCR errors occur in GC rich regions. Ross et al. 14 studied sequencing biases across a large number of genomes. They found that once the homopolymer run is more than 4nt, the insertion and deletion rates start climbing and genomic regions with both high and low GC content are underrepresented in Illumina sequencing (**Figure S1)**. Ananda et al.[25] studied PCR slippage errors and found a rapid increase in homopolymers greater than 4bp. On the other hand, oligos without these characteristics usually exhibit low rates (1%) of synthesis and sequencing errors[6].

**Figure S1:**
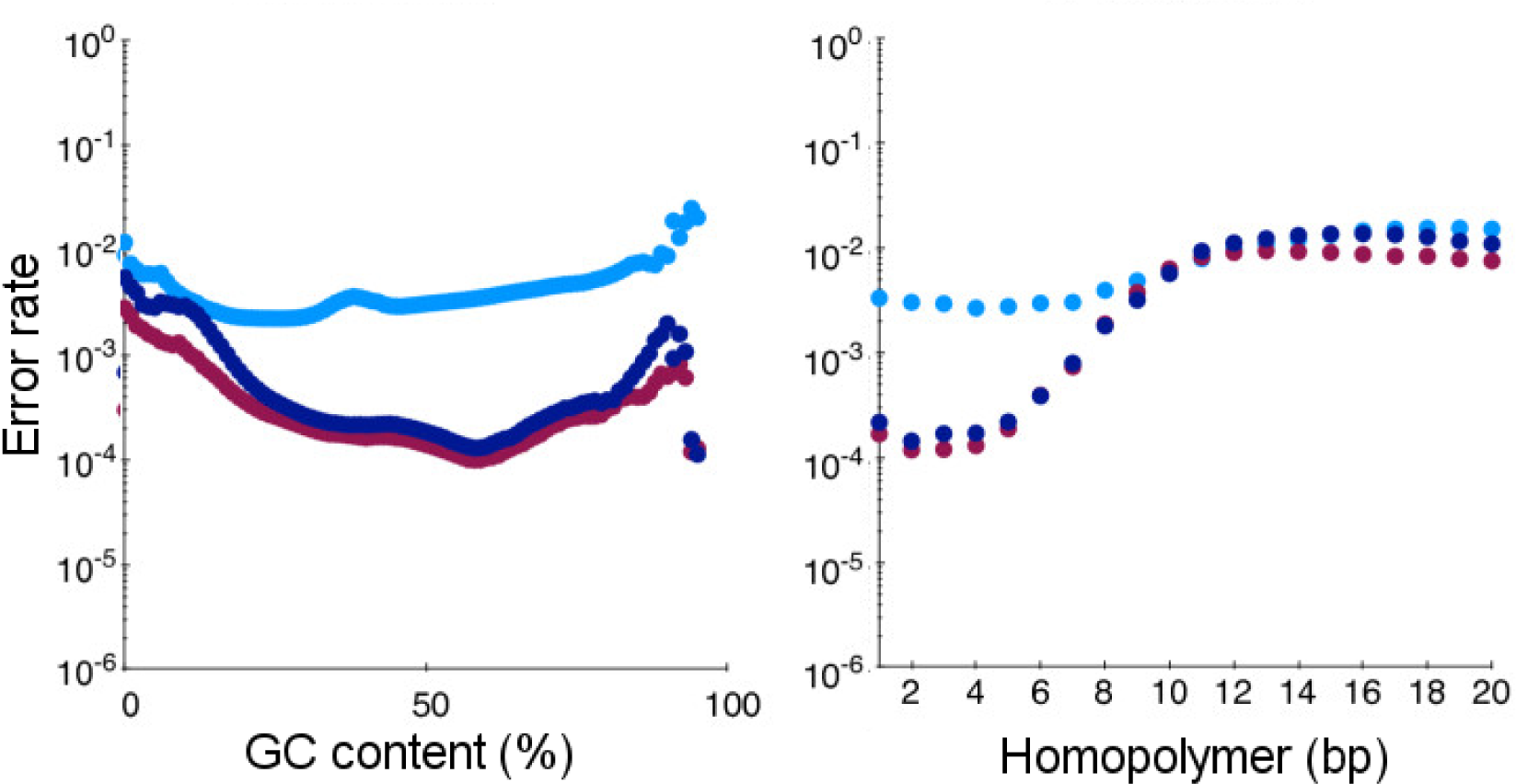
Error rates of Illumina sequencing as a function of GC content and homopolymer length. Light blue: mismatches; dark blue: deletions; purple: insertions. The figure was modified from Ross et al.[14 ] with permission according to publisher license CC-BY-2.0.

To facilitate quantitative analysis, we model the sequence-specific noise by grouping DNA oligos into two classes: valid and invalid. A sequence will be considered valid if its GC content is within 0.5 ±*c*_*gc*_ and its longest homopolymer length is up to m nucleotides. Otherwise, it will be considered invalid and cannot not be transmitted. The coding potential, b, describes the entropy of each nucleotide in valid sequences. Next, valid sequences are exposed to a low *δ*_υ_ dropout rate. Due to the multiplexing architecture of synthesis reactions and high throughput sequencing, the oligos are completely mixed in a pool. Therefore, we need to index each oligo with a short tag. The length of the total oligo will be denoted by l and the fraction of nucleotides that are used for the index will be denoted by i and we will use K to denote the number of segments needed for decoding in the input file. By selecting realistic values for *m, c*_*gc*_, *i*, and *δ*_v_, we can approximate the information capacity, which is defined as the upper bound on the number of bits that can be encoded per nucleotide. Our model does not include synthesis and sequencing errors for valid oligos. Previous work involving DNA storage with high throughput sequencing has shown that it is possible to achieve an error-free consensus for the original oligos by deep sequencing the oligo pool[3, *4, 6*]. We observed a similar pattern in our experiments. When deep sequencing coverage was used (250x) using the master pool, we could reach an error-less recovery without implementing the error correcting code but only a k-mer correction strategy[28]. In a case of low sequencing coverage, our analysis is likely to provide an upper bound on the capacity of DNA storage.

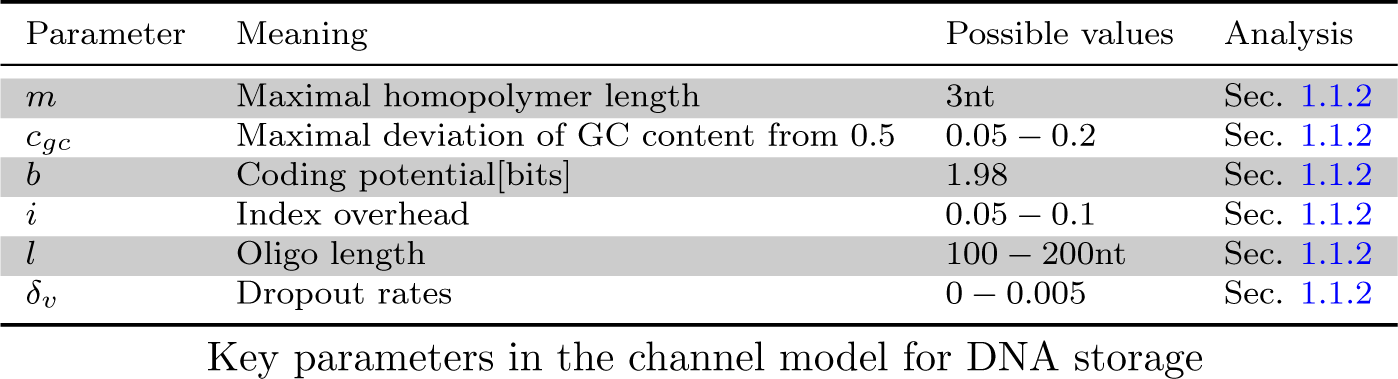
Key parameters in the channel model for DNA storage

| Parameter | Meaning | Possible values | Analysis |
| --- | --- | --- | --- |
| $m$ | Maximal homopolymer length | 3nt | Sec. 1.1.2 |
| $c_{gc}$ | Maximal deviation of GC content from 0.5 | 0.05 – 0.2 | Sec. 1.1.2 |
| $b$ | Coding potential[bits] | 1.98 | Sec. 1.1.2 |
| $i$ | Index overhead | 0.05 – 0.1 | Sec. 1.1.2 |
| $l$ | Oligo length | 100 – 200nt | Sec. 1.1.2 |
| $\delta_v$ | Dropout rates | 0 – 0.005 | Sec. 1.1.2 |

In the next sections, we will show that the information capacity per nucleotide is approximately 1.83bits by setting conservative but realistic parameters.

#### 1.1.2 Quantitative analysis

Under the model above, a DNA storage device behaves as a constrained channel concatenated to an erasure channel (**Figure S2)**

**Figure S2:**
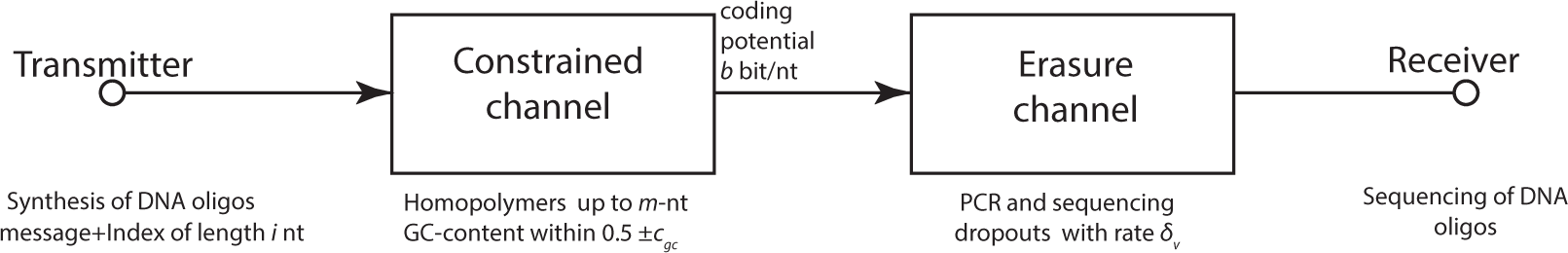
The channel model.

Let *A_X_* be the set of all possible transmitted DNA sequences of length l nucleotides, and *A_Y_* be the set of all possible received sequences. *X* and *Y* denote random DNA sequences from *A_X_* and *A_Y_*, respectively.

The mutual information of *X* and *Y* is defined as:

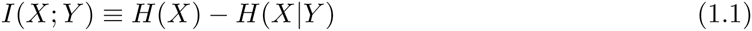

where *H(X)* is the entropy (in bits) of *X* and *H*(*X*|*Y*) is the conditional entropy of *X* given *Y*, or the expected residual uncertainty of the receiver about a transmitted sequence.

The information capacity of each oligo is defined as:

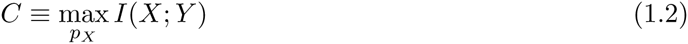

and the information capacity per nucleotide is:

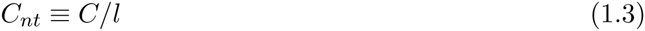

In a constrained channel, the mutual information is maximized by equiprobable transmission of only valid sequences[11 *pp. 250-251*]. Thus, *H*(*X*) = log_2_ |*A_X_*|, where | . | denotes the size of items in a set. Valid sequences are either perfectly received with a probability of 1 – *δ*_*υ*_ or dropped out with a probability of *δ*_υ_. In the former case, the conditional entropy is *H*(*X*|*Y*) = 0, whereas in the latter case the conditional entropy is *H*(*X*|*Y*) = *H*(*X*). Therefore, overall, *H*(*X*|*Y*) = *δ*_υ_*H*(*X*) and the capacity of each nucleotide is:

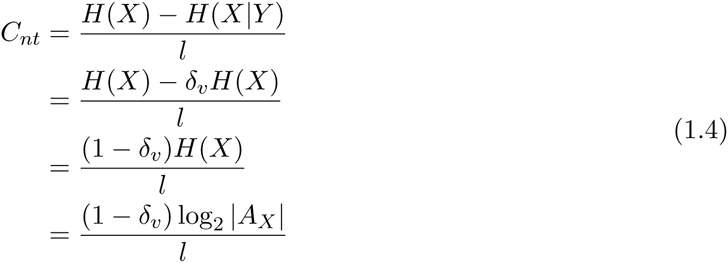

##### The homopolymer constraint

With the homopolymer constraint, the size of the set of all valid code words is:

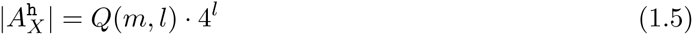

where 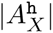 denotes the size of the A_X_ set under the homopolymer constraint and Q(*m,l*) is the probability to observe up to an *m*-nt homopolymer run in a random *l*-nt sequence.

We will start by analyzing a simpler case of the probability of *not* observing a run of m or more successes in *l* Bernoulli trials with a success probability *p* and failure probability of *q* = 1 – *p*, denoted by *q*_*m*_(*p, l*). Feller[*29*] proposed (pp.301-303) a tight approximation for *q*_*m*_(*p, l*):

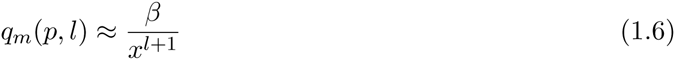

where *x* is:

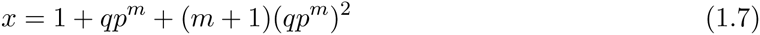

and *β* is:

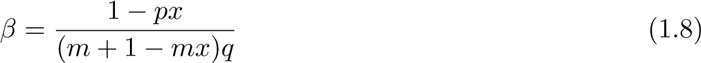

Previous studies have analyzed the general case of the probability distribution function for runs of a collection of symbols from a finite alphabet. The formulae of these distributions are typically defined using recursion or require spectral analysis of matrices with complex patterns [e.g.[*30, 31, 32*] ] and resist analytic analysis. For practical purposes, we approximate the distribution of observing up to m-nt homopolymer runs as the product of four independent events:

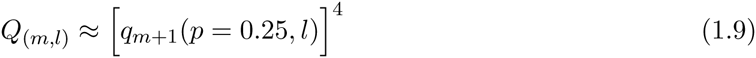

**Table S1** presents the output of Eq. 1.9 versus the expected rate of homopolymers. Overall the approximation is quite consistent with the observed rate for relevant oligo lengths and homopolymer constraints.

**Table S1:** The observed rate of homopolymer mismatches versus the estimated rates from Eq. 1.9

| l | m | Obs. $Q_{(m,l)}$ | Est. $Q_{(m,l)}$ |
| --- | --- | --- | --- |
| 100 | 3 | 0.31 | 0.31 |
| 250 | 3 | 0.05 | 0.05 |
| 100 | 4 | 0.75 | 0.75 |
| 250 | 4 | 0.48 | 0.48 |
| 100 | 5 | 0.94 | 0.93 |
| 250 | 5 | 0.83 | 0.84 |

Combining Eq. 1.5, Eq. 1.6, and Eq. 1.9:

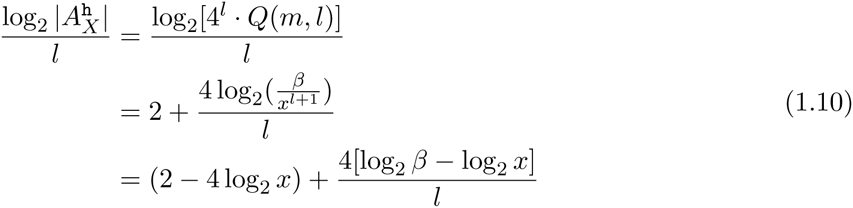

For any *m* **≥** 3 and *l* **≥** 50, we can further approximate:

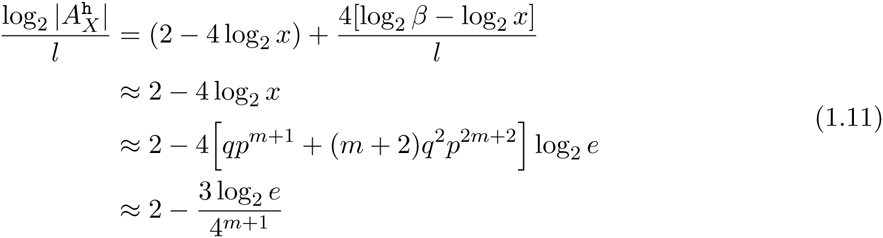

Interestingly, the information capacity per nucleotide under the homopolymer constraint does not depend on the length of the DNA oligos, only on only the maximal length of homopolymers (**Figure S3)**.

**Figure S3:**
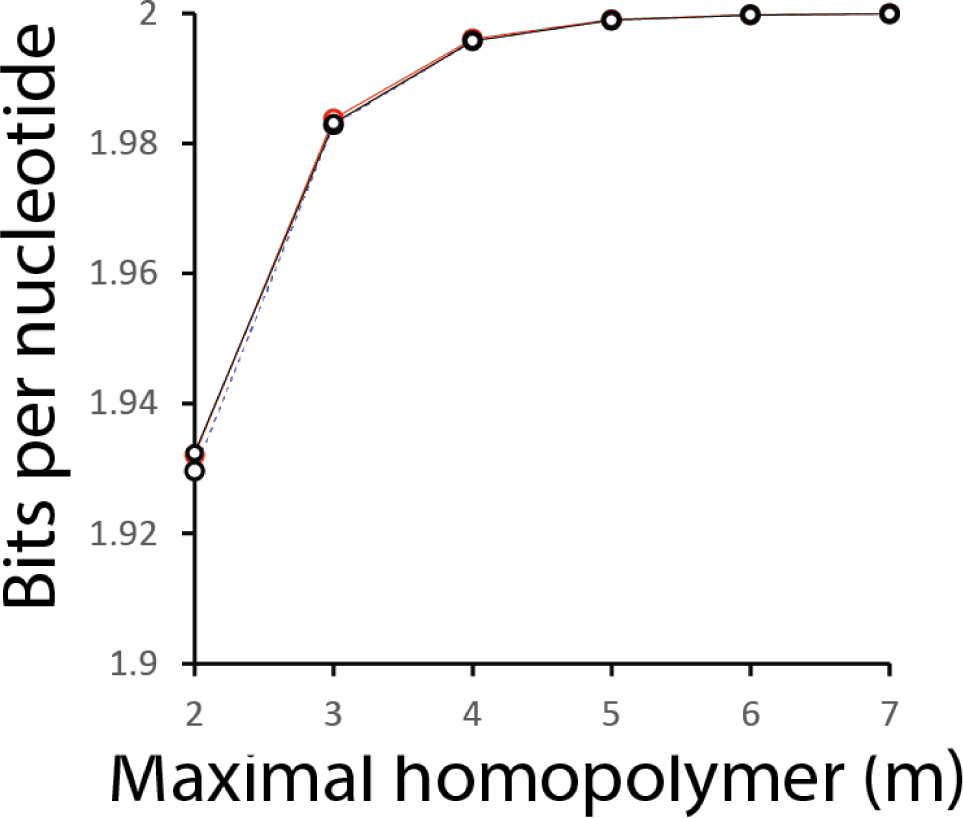
Bits per nucleotide as a function of *m* and *l.* Black: *l* = 50nt, red: *l* = 10^11^nt, broken line: approximating the bits per nucleotide using Eq. 1.11. The three curves almost entirely overlap with each other, illustrating the agreement between Eq. 1.10 and the approximation of Eq. 1.11.

##### The GC content constraint

Let *p*_*gc*_ be the probability that a sequence of *l* nucleotides is within 0.5 ± *c*_*gc*_. Without any other constraint, the *p*_*gc*_(*x*) is:

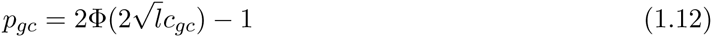

where Φ (†) is the cumulative function of a standard normal distribution. The number of bits per nucleotide that can be transmitted under this constraint is:

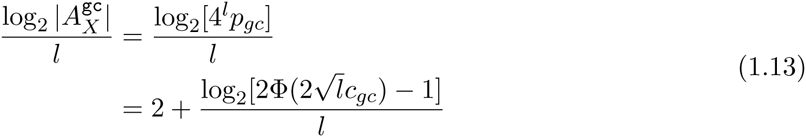

where 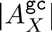 is the size of the *A*_*X*_ set under the GC content constraint.

For any reasonable allowable range of GC content such as *c*_*gc*_ ≥ 0:05 and minimal oligo length of *l* ≥ 50, the GC constraint plays a negligible role in reducing the information content of each nucleotide. For example, with *cgc* = 0:05 and *l* = 100bp, the information content of each nucleotide is 1:992bits, only 0:4% from the theoretical maximum. **Figure S4** presents the channel capacity as a function of the oligo length under various levels of GC content constraints.

**Figure S4:**
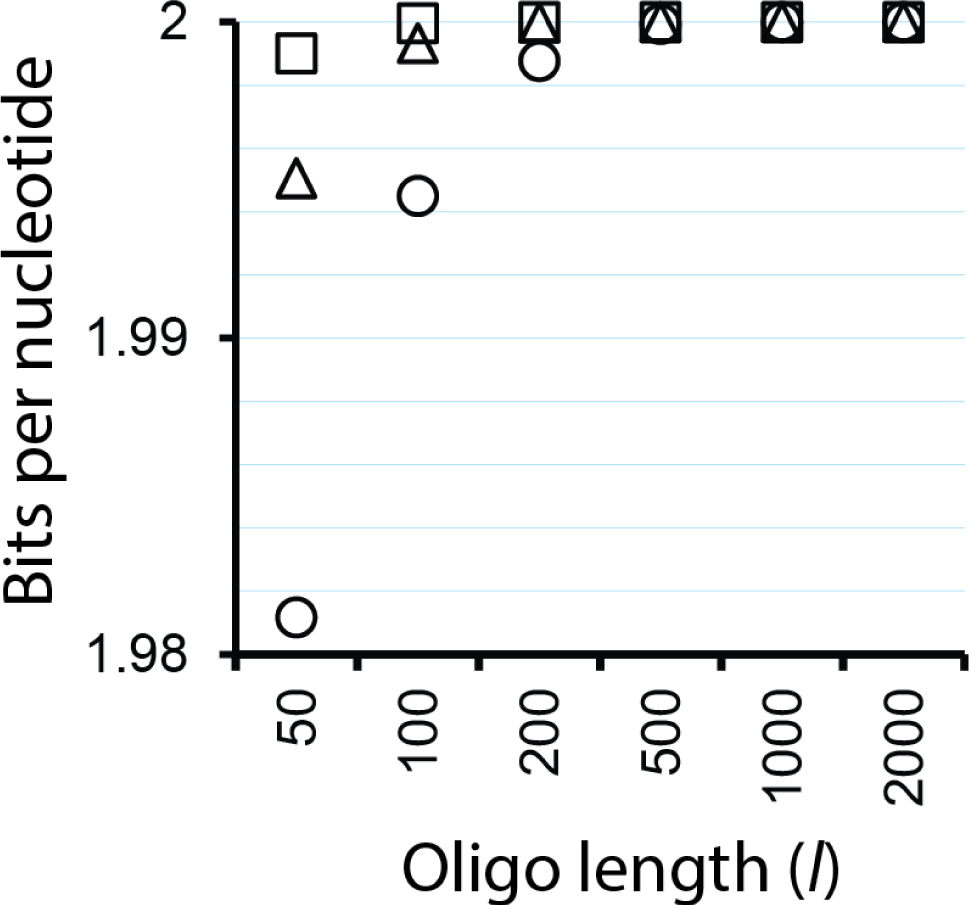
Bits per nucleotide as a function of *l* and *c*_*gc*_. Circles, triangles, and squares denote *c*_*gc*_ = (0.05, 0.1,0.15), respectively.

##### Putting the biochemical constraints together

The homopolymer constraint and the GC content constraint define the output of the constrained channel and *b*, the coding potential per nucleotide.

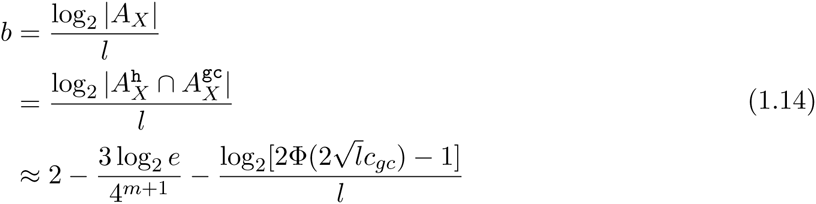

To estimate *b*, we selected a conservative yet practical set of constraints with *c*_*gc*_ = 0:05 and *m* = 3, and an oligo length of *l* = 150nt. We selected *m* = 3 following the work of [*3, 25, 14*] that studied the rates of homopolymer errors in synthesis, PCR ampli_cation, and sequencing, respectively. For the GC content, we decided to use a conservative value of *c*_*gc*_ = 0:05 since this constraint hardly changes the results. *l* = 150 was set to match our experiment and is close to the maximal oligo lengths of major manufacturers before adding two 20nt annealing sites for PCR primers. In this setting, the coding potential is:

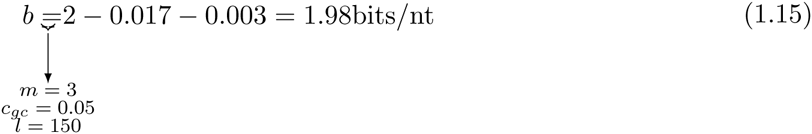

##### Index length

Each oligo should be indexed since they are not received in any particular order. This requirement means that some of the nucleotides in the oligo cannot be used for encoding DNA information. Let *l*‘ denote the remaining nucleotides available for encoding data. Then,

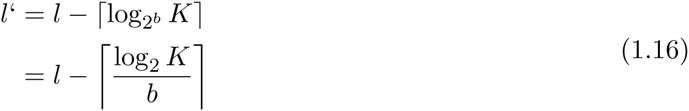

The information capacity of each nucleotide after adding the index is reduced to:

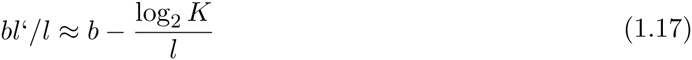

When the oligo length is fixed, it is easy to see that the information capacity goes to zero when the input file becomes larger since indexing would eventually occupy all real estate available on the oligo. However, this issue can be solved by first splitting the file into large blocks of information and encoding each block using a distinct oligo pool that is physically isolated from the other pools, similar to physical hard-drives in an external storage device. For example, we can envision an extreme architecture of a DNA storage architecture that consists of an array of 16Tbytes per oligo pool (comparable to the largest single hard-drive available today). With 150nt oligos, indexing would take only 20nt. This translates to a 13% reduction of the effective oligo length, meaning an information capacity of 1.77bits/nt. In a more practical architecture of an array of oligo pools of 1Mbyte to 1Gbyte of data, the indexing cost goes down to 8 to 13 nucleotides, respectively. This would translate to approximately 7% reduction in the coding capacity to *C_nt_* = 1.84bits/nt assuming no dropouts.

##### Dropouts

Finally, we have to consider the probability of oligo dropouts on the channel capacity. Previous studies have found that sequencing coverage follows a negative binomial distribution (NB) with average of *µ* and a size parameter *r*. The average coverage *µ* is largely determined by the capacity of the sequencer, whereas *r* corresponds to the library preparation technology. When r goes to infinity, then the distribution behaves as Poisson. In practice, however, r is usually between 2 and 7. For example, Sampson et al.[*33*] found that *r* = 2 in an exome sequencing dataset and *r* = 4 for whole genome sequencing using Illumina. In our experiments, we observed *r* = 6.4 for the master pool that used a relatively small number of PCR rounds and *r* = 1.3 for the deep copy that underwent 90 PCR cycles. In the main text, we report the overdispresion, which is defined as 1/r, because it it more intuitive to understand that larger values are more overdispresed.

To find the expected rate of dropouts, we need to evaluate the probability of getting zero sequencing coverage. Let *P*(*x; µ, r*) be the probability mass function of a negative binomial distribution. In general:

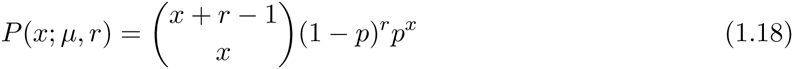

where *p* = *r*/(*r* + *µ*). Then, the rate of dropouts is

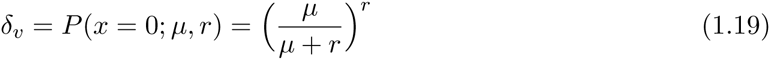

We found an excellent agreement between the model and the empirical number of dropouts. For example, in the downsampling experiments, we reduced the average coverage per oligo to *µ* = 5.86 (the size parameter is invariant to downsampling and stayed at *r* = 6.4). Eq. 1.19 predicted a dropout rate of 1.5%, whereas the observed rate of missing oligos was δ*_v_* = 1.3%. **Figure S5** shows the expected dropout rates for a range of average sequencing coverage and as a function of the size parameter for the relevant range of oligo experiments.

**Figure S5:**
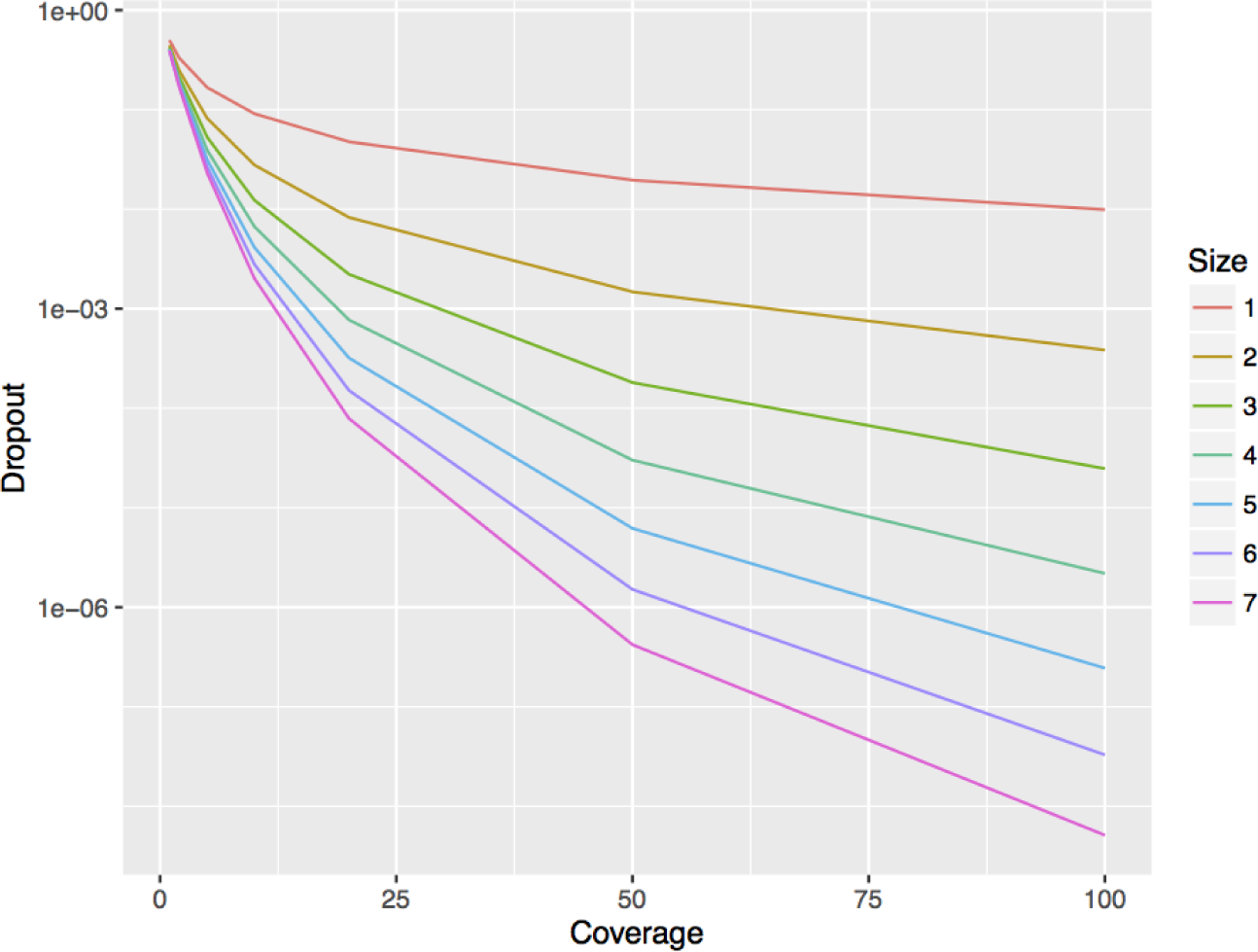
The expected dropout rate *(δ*_υ_) as a function of the average sequencing coverage (*µ*) and the size parameter *r*.

**Figure S6:**
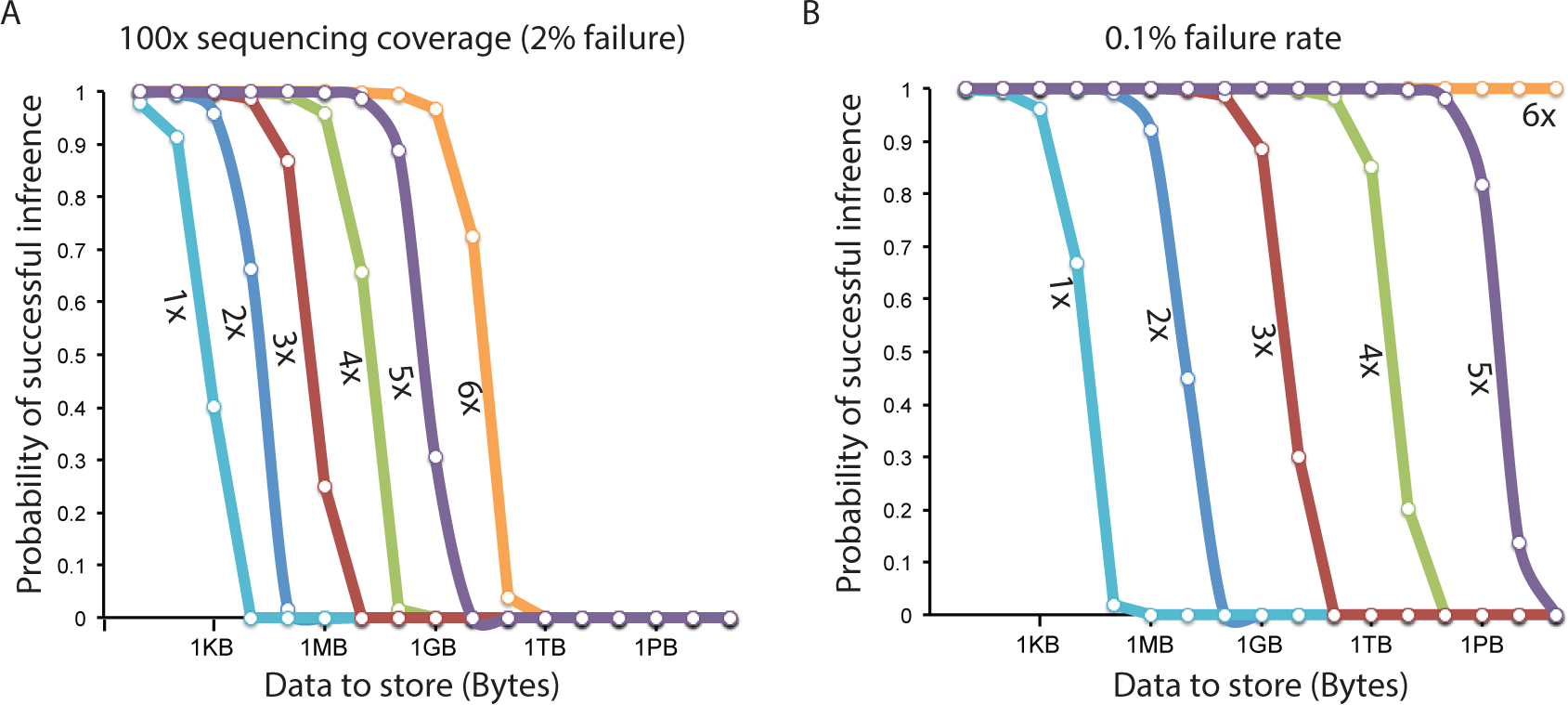
The probability of perfect retrieval (no gaps) of information as a function of input size and the number of times each input bit is repeated on distinct oligos (colored lines) (A) Dropout rate of 2%. This rate was observed in Goldman et al.[*4*] after downsampling their sequencing data to reflect a coverage of 100 × (B) Dropout rate of 0.1%, nine times smaller than the rate observed by Blawat et al.[*7*]. With this low drop out rate, 1Gbyte storage has 10% chance of data corruption even when each bit is stored on 3 distinct oligos.

These results show that for a reasonable sequencing coverage of *µ* = 10 and a relatively mediocre size parameter of *r* = 2, the expected dropout rate is 2.7%. When the size parameter is better with *r* = 6, the dropout rate is approximately 0.25%. We posit that in most storage architectures, the dropout rates should be around 5 = 0.5%. When the size parameter is excellent (*r* = 7), this dropout rate can be achieved with a low coverage of *µ* = 7. When the size parameter is of low quality (*r* = 1.5), one can achieve this rate with sequencing coverage *µ* = 50, which is not unreasonable even for large oligo pools. Blawat et al.[7] recently reported an average dropout rate of 0.9% in an experiment with four batches of quarter million oligos before employing an aggressive correction of short oligos using their error correcting code, close to the rate in our analysis.

According to Eq. 1.3, the reduction in capacity per nucleotide is proportional to the dropout rate. In the previous section, we found that the capacity is *C*_*nt*_ = 1.84bits/nt without any dropout. With a dropout rate of 0.5%, the capacity is *C*_*nt*_ = 1.83bits/nucleotide.

### 1.2 Calculating the results of DNA storage schemes

1. **Church et al.**[*3*] used a binary scheme with a coding potential of *b* = 1bit/nt. The scheme has no error correcting code or fold redundancy. The index is of length 19bits. They encoded 658, 750bytes using 54, 898 oligos of length 115nt. The net information density is 658, 750 x 8/(54,898 x 115) = 0.83bits per nucleotide. The supplemental material of the Church et al. experiment states that 10 femtomole of DNA were used to retrieve the data. This gives a total of 10 × 10^−15^ 6.02 × 10^23^ = 6.02 × 10^9^ DNA molecules. The physical density is therefore 658, 750[bytes]/(6.02 × 10^9^[molecules] × 159[nt/molecule] × 325[Dalton/nt] × 1.67 × 10^−24^[g/Dalton]) = 1.28 × 10^15^ [byte/g].
2. **Goldman et al.**[4] use a ternary scheme with *b* = 1.58bit/nt. The scheme has one parity nucleotide for error detection and two nucleotides to detect reverse complement coding and a 4 fold redundancy. The index is of length 14trits, 2trits for file locations and 12 trits for an address, which is equivalent to a 21bit index. They encoded 757, 051bytes using 153,335 oligos of length 117nt. The net information density is 757051 × 8/(153, 335 × 117) = 0.34bits per nucleotide. The supplemental material of the Goldman et al. experiment states that they used 337pg of input DNA to retrieve the information. The physical density is 757051[bytes] /(337×10^−12^[g]) = 2.25 × 10^15^[byte/g].
3. **Grass et al.**[*5*] use a GF(47) scheme that maps every two bytes to nine nucleotides with b = 1.78bit/nt. The scheme uses a 3byte index and a two dimensional error correcting code that mapped 30 blocks of data of lengths 594bytes to an array of 713 blocks of size 39bytes including a 3byte index. They used two dimensional Reed-Solomon codes to correct dropouts. They encoded 82, 896bytes using 4, 991 oligos of length 117nt. The net information density is 82896 × 8/(4, 991 × 117) = 1.14bits per nucleotide. Grass et al. stated that they used appro×imately 1ng. The weight per molecule is 158[nt/molecule] ×320[Dalton/nt] ×1.67 × 10^−24^[g/Dalton] = 8.44 × 10^−20^[g/molecule]. The total number of molecules in the reaction is 10^−9^/(8.44 × 10^−20^) = 1.18 × 10^10^ DNA molecules. With 4991 oligos, the average number of molecules per sequence is 2,370,000 copies. The physical density is 4991[bytes]/1 × 10^−9^[g] =5 × 10^12^[byte/g].
4. **Bornholt et al.**[*6*] used the same scheme as Goldman et al. but with 1.5 fold redundancy against dropouts. The length of the index is not mentioned in the manuscript. They encoded 45,652bytes and synthesized 151,000 oligos of length 120, but their experiments included testing the original Goldman et al. scheme for their data. We therefore estimated the net density by reducing the redundancy of Goldman et al. from 4× to 1.5×. With this estimate, the net information density is 0.34 × 4/1.5 = 0.88bits per nucleotide. We were unable to determine the physical density of the study due to lack of details.
5. **Blawat et al.**[*7*] use a scheme that maps each input byte to 5 nucleotides and achieves b = 1.6bits/nt. The index is of length 39bits and the scheme uses a two dimensional nested error correcting code. First, the index is mapped to a 63bit vector. Next, the data and the 63bit index are organized into a two dimensional array. The scheme employs a Reed Solomon code that adds 33bits of redundancy to each block of 223bits in one dimension and 16bits of cyclic redundancy check in the other dimension. They encoded 22MByte of data using 1,000,000 oligos of length 190nt and showed that their scheme can decode dropouts. The net information density is 22 × 10^6^ × 8/(10^6^ × 190) = 0.92bit per nucleotide. We were unable to determine the physical density of the study due to lack of details.
6. **This work** uses a scheme that screens potential oligos to realize the maximal coding capacity with *b* = 1.98bit/nt. The seed has 4bytes that can encode files of up to 500MByte (see section 1.3.5). We also add 2byte of Reed-Solomon error correcting code that protects both the seed and the data payload. The level of redundancy against dropouts is determined by the user and we decided on a level of 1.07× because of the proposed cost from the oligo manufacturer. We encoded 2,146, 816bytes of information using 72, 000 oligos of length 152nt. The net information density is 2146816 × 8/(72000 × 152) = 1.57bits per nucleotide. The physical density is discussed in the main text.

We are fully aware that some of the differences in the net information densities can be attributed to different oligo lengths or indexes. We decided to present the net density without standardizing the schemes to a specific oligo/index length for several reasons: first, certain schemes, such as Blawa et al.[*7*], employ error correcting codes that are designated for specific input and output lengths. It is not ea sy to translate the scheme to a different oligo length without completely changing their error correcting strategy. Second, our main focus is to compare schemes that were tested in practice. Imputing the net density for a different architecture without empirical tests can be misleading. For example, Goldman et al.[*4*] reported a successful filtering of sequencing errors even with low sequencing coverage using a single parity nucleotide when the length of oligos was 117nt. It is not clear whether this error detection scheme can work with 150nt oligos that are more error prone. Third, we found that the standardization does not significantly affect the overall picture. For example, after standardizing the Church et al. scheme to have an index length of 28bits and oligo length of 152nt (similar to our method), the net density goes from 0.83bit/nt to 0.81bit/nt. Similarly, Goldman et al. goes from 0.34bit/nt to 0.28bit/nt after standardization.

Throughput the manuscript, we used the scientific SI units to indicate the scale of digital information and not the Joint Electron Device Engineering Council (JEDEC) standard that uses the binary system. For example, 1Mbyte refers to 10^6^ bytes and not 2^20^ bytes and 1Pbyte refer to 10^15^ bytes and not 2^50^ bytes.

The physical density was calculated including the adapter regions for compatibility with the Church et al. study that pioneered this calculation.

### 1.3 The DNA Fountain coding strategy

Our encoding algorithm works in three computational steps: (a) preprocessing, (b) Luby Transform, and (c) screening. Its overall aim is to convert input files into a collection of valid DNA oligos that pass the biochemical constraints and be sent to synthesis.

#### 1.3.1 Encoding overview

1. In the preprocessing step, we start by packaging the files of interest into a single tape-archive (tar) file, which is than compressed using a standard lossless algorithm (e.g. gzip). Besides the obvious advantage of reducing the size of the tar file, compression increases the entropy of each bit of the input file and reduces local correlations, which is important for the screening step. Then, the algorithm logically partitions the compressed file into non-overlapping segments of length *L* bits, which is a user defined parameter. We used *L* = 256bits (32 bytes) for our experiments, since this number is compatible with standard computing environments and generates oligos of lengths that are within the limit of the manufacturer.
2. The Luby Transform step works as follows:

a. We initialize a pseudorandom number generator (PRNG) with a seed, which is selected according to a mathematical rule as explained in Section 1.3.2.
b. The algorithm decides on *d*, the number of segments to package in the droplet. For this, the algorithm uses the PRNG to draw a random number from a special distribution function, called robust soliton probability distribution. Briefly, the robust soliton distribution function is bi-modal and ensures that most of the droplets are created with either a small number of input segments or a fixed intermediary number of segments. This mathematical property is critical for the decoding process. Section 1.3.3 presents this distribution in details.
c. The algorithm again uses the PRNG to draw *d* segments without replacement from the collection of segments using a uniform distribution.
d. The algorithm performs a bitwise-XOR operation (biwise addition modulo 2) on the segments. For example, consider that the algorithm randomly selected three input fragments: 0100, 1100, 1001; In this case, the droplet is: 0100 ⊕ 1100 ⊕ 1001 = 0001.
e. The algorithm attaches a fixed-length index that specifies the binary representation of the seed. For example, if the seed is 3 and the fixed index length is 2bits, the output droplet will be 110001. In practice, we used a 32bit (4byte) index for compatibility with standard computing environments.
f. the user has the option to use a regular error correcting code computed on the entire droplet. In our experiments, we added two bytes of Reed-Solomon over *GF*(256) to increase the robustness of our design. The Luby Transform confers robustness against dropouts. Theoretically, the transform additions can be thought of as representing the input segments as a binary system of linear equations. Each droplet is one equation, where the seed region has one to one correspondence to the 0-1 coefficients of the equation, the payload region is the observation, and the data in the input segments are the unknown variables of the system. To successfully restore the file, the decoder basically needs to solve the linear system. This task can theoretically be done (with a very high probability) by observing any subset of a little more than *K* droplets. Our encoder exploits this property to create robustness against dropouts. It produces many more oligos than the dropout rates (eg. %5 more oligos) to create an over-determined system. Then, no matter which oligos are dropped, we can solve the system and recover the file as long as we can collect a little more than *K*. We postpone the formal definition of ”a little more” to Section 1.3.3 that presents the robust solition distribution.
3. In the screening step, the algorithm excludes droplets that violate the required biochemical constraints from the DNA sequence. First, it converts the droplet into a DNA sequence by translating {00,01,10,11} to {A,C,G,T}, respectively. For example, the droplet “110001” corresponds to “TAC”. Next, the algorithm screens the sequence for desired properties such as GC content and homopolymer runs. If the sequence passes the screen, it is considered valid and added to the oligo design file; otherwise, the algorithm simply trashes the sequence. Since the compressed input file essentially behaves as a random sequence of bits, screening has no effect on the distribution of the number of source fragments in the valid droplets. For example, a droplet that was created by XOR-ing 10 source fragments has the same chance of passing the screen as a droplet with only 1 source fragment. This asserts that the number of droplets in a valid oligo follows the robust soliton distribution regardless of the screening, which is crucial for the decoder.

We continue the oligo creation and screening until a desired number of oligos are produced. The decoder outputs a FASTA file that can be sent to the oligo manufacturer.

#### 1.3.2 Seed schedule

The process of droplet creation starts with a seed to initialize the pseudorandom number generator. Typical fountain code implementation starts with a specific seed that is incremented with every round. The problem in our case is that an incremental seed schedule creates bursts of invalid sequences. For example, any seed in interval [0,1,…, 16777215] would be mapped to a sequence that starts with an `AAAA` homopolymer when representing the seed as a 32bit number.

We sought a strategy that would go over each number in the interval [1,…, 2^32^–1] in a random order to avoid bursts of invalid seeds. For this we used a Galois linear-feedback shift register (LFSR). In this procedure, a hard-coded seed (e.g. 42) is represented as a binary number in the LFSR. Then, we deploy one round of the LFSR, which shifts and performs a XOR operation on specific input bits within the register. This operation corresponds to a polynomial multiplication over a finite field. The new number in the register is used as the seed for the next droplet. By repeating this procedure, we can generate a sequence of seeds in a pseudo-random order. To scan all numbers in the interval, we instructed the LFSR to use the primitive polynomial *x*^32^ + *x*^30^ + *x*^26^ + *x*^25^ + 1 for the multiplication of the number in the register. With this polynomial, the LFSR is guaranteed to examine each seed in the interval [1,…, 2^32^ – 1] without repetition. Other implementations of DNA Fountain might require a different seed space. This can easily be done by switching the polynomial in the LFSR. Public tables such *as[34*] list LFSR polynomials with a cycle of *2*^*n*^ – 1 for a wide range of *n*.

#### 1.3.3 Tuning the Soliton distribution parameters

The robust soliton distribution function, *µ*_*k,c*,δ_ (d) is a key component of the Luby Transform. We will start by describing two probability distribution functions, ρ(*d*) and τ(*d*), that are the building blocks for the robust soliton distribution function.

Let ρ(*d*) be a probability distribution function with:

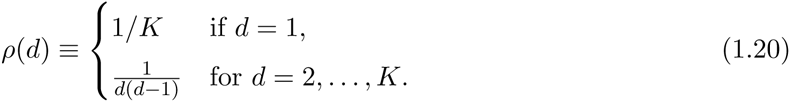

Let τ be a probability distribution function with:

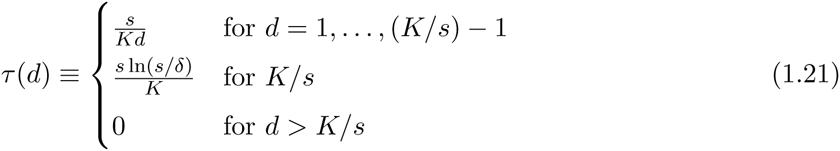

where 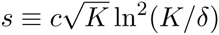.

The robust soliton distribution is defined as:

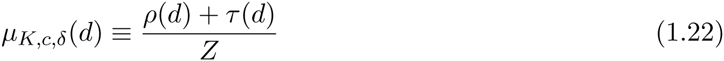

Where *Z* is a normalization parameter, *Z* = Σ_*d*_ ρ(*d*) + t(*d*) = 1.

**Figure S7** presents an example of the robust soliton distribution.

**Figure S7:**
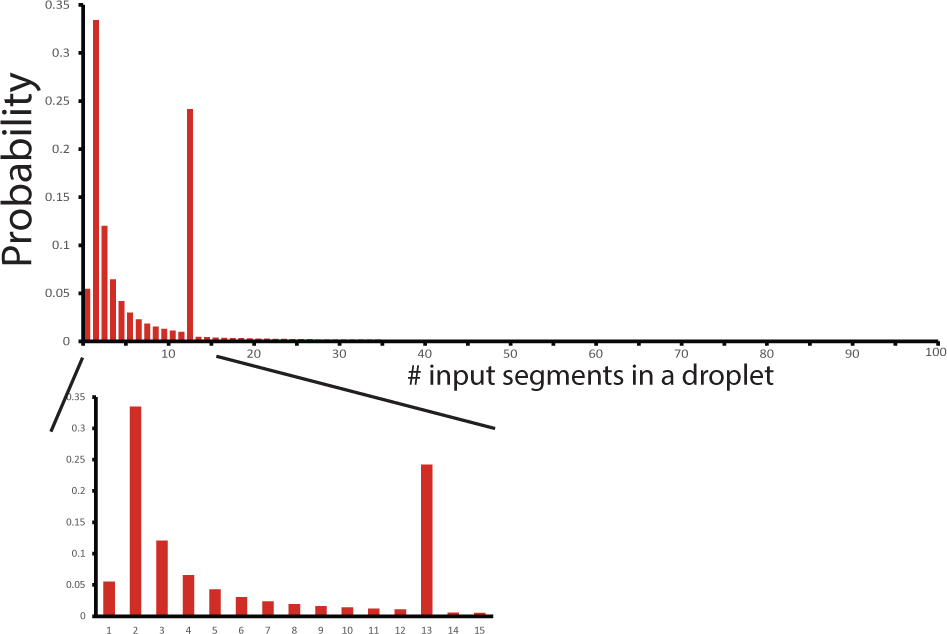
An example of the Soliton distribution with (*K* = 100, *c* = 0.1, *δ* = 0.05). Most droplets will include 1-3 input segments and a large fraction of the remainder will include 13 segments. The c and *δ* parameters are controlled by the user.

The two free parameters, c and 5, are specified by the user during the oligo synthesis stage. In general, 5 is the upper bound probability of failing to recover the original file after receiving *K* · *Z* droplets[*18, 19*]. Previous mathematical analyses have shown that this upper bound is highly pessimistic and that the probability of a failure is much smaller in practice[*19*]. The *c* parameter is a positive number that affects the decoding/encoding performance. On one hand, low values of *c* reduce the *average* number of oligos that are required for successful decoding. On the other hand, low values of *c* increase the *variance* around the average[*19*]. In addition, low values of *c* increase the average number of segments in each droplet. This translates to deeper recursions of the decoding algorithm and increases the runtime[*35*].

In our experiments, we selected *δ* = 0.001 and *c* = 0.025. With these values, *Z* = 1.033, meaning that we needed to generate at least 3% more oligos than the number of segments.

#### 1.3.4 Decoding

We employed the following steps to decode the data:

1. Preprocessing: first, we stitched the paired-end reads using PEAR[36] and retained only sequences whose length was 152nt. Next, we collapsed identical sequences and stored the collapsed sequence and its number of occurrences in the data. Finally, we sorted the sequences based on their abundance so that more prevalent sequences appear first. This way the decoder observes the highest quality data first and gradually gets sequences with reduced quality. Due to the redundancy of our approach, the decoder usually does not need all oligos to construct the original file and will usually stop before attempting to decode sequences that were observed a small number of times (e.g. singletons), that are more likely to be erroneous. From this point the decoder process works sequentially and executes the next two steps on each collapsed sequence until the file is fully resolved:
2. Droplet recovery: the decoder maps the DNA sequence into a binary format by translating {A,C,G,T}, to {0; 1; 2; 3}. Next, it parses the sequence read to extract the seed, data payload, and the error correcting code (if it exists). If there is an error correcting code, the decoder checks whether there are any errors. In our experiments, we excluded sequences with an indication of one or more errors and did not attempt to correct them. We found that most errors were due to short insertions and deletions, potentially from the oligo synthesis. Reed-Solomon error correcting code can only handle substitutions and attempting to correct the sequence is more likely to result in erroneous recovery. In addition, the user can instruct the decoder also for an additional error detection approach using *k*-mers. Under this option, the decoder will first compare a new sequence read to all previously processed reads that were used for decoding. If there is at least one match of a stretch of *k* nucleotides than the new sequence would be discarded. The idea is that erroneous reads should show a high but not perfect similarity to valid oligos that were observed before.
3. Segment inference: after validating the integrity of the binary message, the decoder initializes a pseudorandom number generator with the observed seed. This generates a list of input segment identifiers. Next, it employs one round of a message passing algorithm, which works as follows: first, if the droplet contains segments that were already inferred, the algorithm will XOR these segments from the droplet and remove them from the identity list of the droplet. Second, if the droplet has only one segment left in the list, the algorithm will set the segment to the droplet’s data payload. Next, the algorithm will propagate the information about the new inferred segment to all previous droplets and will repeat the same procedure recursively, until no more updates can be made. If the file is not recovered, the decoder will move to the next sequence in the file and execute the droplet recovery and segment inference steps. This process of propagation of information eventually escalates that solves the entire file (**Figure S10**).

#### 1.3.5 DNA Fountain overhead

##### Code rate

DNA Fountain approaches the channel capacity but still entails a small overhead for a wide range of realistic applications. First, even with zero probability of dropout, we need to synthesize more oligos than the number of input segments. As discussed in section 1.3.3, the robust soliton distribution asserts convergence of the decoder (with a probability of 5) when *K* · *Z* droplets are seen. In general, when *K* is extremely large, *Z* → 1, meaning that the code is rateless and entails no overhead. However, for experiments with tens of thousands of oligos, *Z* is on the order of 3% and empirical results with 10000 segments have observed on average a 5% overhead for successful decoding[*37*]. Second, our strategy entails a small inflation in the size of the index. Eq. 1.16 shows that the minimal index length is log_2_*b K* = log_3.95_ *K*. In our case, the index space must be larger to accommodate both successful and failed attempts to construct a droplet. This inflation is quite modest and only scales logarithmically with the size of the search space. For example, when screening 150nt oligos for a GC content between 45% and 55% and homopolymer runs of up to 3nt, only 12.5% of the oligos pass these conditions. This requires the index to have an additional ⌈log_3.95_(1/0.125)⌉ = 2 nucleotides, which reduces the information content by 2/150=1.3% from the theoretical maximum. Thus, we posit that our approach could theoretically reach 100% – (3% + 1.3%) ≈ 96% of the channel capacity for experiments with tens of thousands of oligos.

In practice, we realized 1.57/1.83 = 86% of the channel capacity. The difference between the practical code rate and the theoretical rate is explained by three factors. First, our coding strategy included a redundancy level of 7%, about 3% more than the required overhead for successful decoding with 0.5% dropout rate. As explained in the main text, we selected this level mainly because of the price structure of the oligo manufacturer that offered the same price for any number of oligos between 66,000 to 72,000. We consider the flexibility of DNA Fountain to maximize the number of informative oligos within a specific price range as a strong advantage of our technique. For the seed space, we used 32bits. This seed is about 9bits over what is needed for encoding all successful and failed attempts to create valid oligos for a file of size 2.1Mbyte. We decided to use this length in order to be able to scale the same architecture to store files of 500Mbyte. Finally, we also had an error correcting code of two bytes in each oligo, which reduced the rate by another 5%.

##### The time complexity and economy of encoding and decoding

Our experiments show that DNA Fountain is feasible for a range of file sizes and oligo lengths. From a computation complexity perspective, our encoding strategy scales log-linearly with the input file size and empirical tests showed that a 50Mbyte file takes about 1 hour on a single CPU of a standard laptop. On the other hand, the complexity scales exponentially with the oligo length because it becomes harder to find sequences without homopolymers and increasing numbers of droplets are created only to be destroyed in the screening process. However, the base of the exponent is very close to 1. For example, with *m* = 3, the complexity scales with 𝒪(1.01^*l*^) and with *m* = 4 the complexity scales with 𝒪(1.003^*l*^). With this low exponent we found that our strategy is still practical for a range of oligo lengths even above the limit of major oligo pool manufacturers of approximately 200-250nt[*22*]. For example, encoding data on 300nt oligos takes only 3min for 1Mbyte per CPU and encoding 400nt oligos takes 6min for the same conditions (**Table S2**).

**Table S2:** Encoding time and cost saving using DNA Fountain. Run time indicates the empirical time it takes the encoder to convert an input file to oligos for one CPU. Computing costs show an estimate of the price to encode the file using Amazon Cloud with a price of $0.5 per hour for a 40 CPU server. Oligo synthesis money saved reports the estimated difference in budget between our technique and previous techniques with an information capacity of approximately 0.9bit/nt. We assume a constant price for different oligo lengths and sizes of $640 per 1Mbase. Saving was rounded to the closet thousand. The upper part of the table reports the results for different input file sizes and an oligo length of 152nt. The bottom part reports the results for different oligo lengths and a 1Mbyte input file.

| | Condition | Run time[sec] | Computing costs [cents] | Oligo synthesis money saved [\$] |
| --- | --- | --- | --- | --- |
| Input file | 1Mbyte | 50 | 0.02 | 5,000 |
|  | 2Mbyte | 112 | 0.04 | 11,000 |
|  | 4Mbyte | 200 | 0.07 | 23,000 |
|  | 8Mbyte | 436 | 0.15 | 45,000 |
|  | 16Mbyte | 897 | 0.31 | 96,000 |
|  | 32Mbyte | 1817 | 0.63 | 192,000 |
|  | 64Mbyte | 3699 | 1.28 | 377,000 |
| Oligo length | 100nt | 53 | 0.02 | 5,000 |
|  | 152nt | 40 | 0.01 |  |
|  | 200nt | 54 | 0.02 |  |
|  | 252nt | 84 | 0.03 |  |
|  | 300nt | 137 | 0.05 |  |
|  | 352nt | 222 | 0.07 |  |
|  | 400nt | 374 | 0.13 |  |

**Table S3:** Summary of the dilution experiment results. ^(1)^ total molecules were calculated based on dividing the total weight by the expected weight for 200nt single-strand DNA: 1.0688 × 10^−^19 gr/molecule. ^(2)^ The number of molecules divided by 72000, the total number of synthesized oligos. ^(3)^ The fraction of PE reads that were assembled by PEAR. ^(4)^ The average number of reads per valid oligo. ^(5)^ The size parameter (overdispersion^−^1) of the negative bionomial distribution. o.o.w: out of which.

| **Dilution** | **1** | **2** | **3** | **4** | **5** | **6** | **7** |
| --- | --- | --- | --- | --- | --- | --- | --- |
| Total weight[ng] | 10 | 1 | 0.1 | 0.01 | 0.001 | 0.0001 | 0.00001 |
| Total molecules<sup>(1)</sup> | $9.21 \times 10^{10}$ | $9.21 \times 10^9$ | $9.21 \times 10^8$ | $9.21 \times 10^7$ | $9.21 \times 10^6$ | $9.21 \times 10^5$ | $9.21 \times 10^4$ |
| Molecules per sequence<sup>(2)</sup> | $1.3 \times 10^6$ | $1.3 \times 10^5$ | $1.3 \times 10^4$ | $1.3 \times 10^3$ | $1.3 \times 10^2$ | $1.3 \times 10$ | 1.3 |
| Density[byte/g] | $2.14 \times 10^{14}$ | $2.14 \times 10^{15}$ | $2.14 \times 10^{16}$ | $2.14 \times 10^{17}$ | $2.14 \times 10^{18}$ | $2.14 \times 10^{19}$ | $2.14 \times 10^{20}$ |
| Total reads | 40,843,415 | 76,865,956 | 66,855,708 | 65,706,638 | 55,996,408 | 62,050,438 | 63,510,382 |
| ... Reads assembled(%)<sup>(3)</sup> | 93 | 95 | 97 | 94 | 94 | 94 | 84 |
| .....o.o.w. length 152nt(%) | 66 | 67 | 70 | 67 | 68 | 69 | 65 |
| .....o.o.w valid oligos(%) | 89 | 90 | 89 | 89 | 90 | 90 | 90 |
| Reads passed all stages(%) | 45 | 43 | 40 | 44 | 42 | 42 | 50 |
| Recovered oligos | 72000 | 72000 | 71998 | 69407 | 27379 | 14194 | 9651 |
| Missing oligos (%) | 0 | 0 | 0 | 4 | 62 | 80 | 87 |
| Mean coverage<sup>(4)</sup> | 311 | 611 | 559 | 511 | 448 | 501 | 439 |
| Size<sup>(5)</sup> | 7.49 | 7.46 | 4.33 | 0.97 | 0.06 | 0.02 | 0.01 |

Our ability to include more information in each nucleotide compensates for the computation price even for long oligos. For example, encoding 1Gbyte of information on 400nt oligos would take 4 CPU days. We can parallelize the encoder on multiple CPUs by letting each encoder thread scan a different region in the seed space. With the current price of $0.5 for one hour of a 40 CPU server on Amazon Web Services, we can encode the entire input data for $1.25 in a few hours. The price of the computing time is substantially smaller than the cost reduction in the oligo synthesis costs. Assuming that oligo synthesis is $640 per 1Mbase (current price for our experiment was $7000 and synthesized 152 x 72000 nucleotides without Illumina adapters), synthesizing 1Gbyte with our scheme would cost approximately $3.27 million. However, when using other methods with lower information content of 0.9bit/nt, the synthesis price is nearly $5.63 million, rendering the computational costs marginals compared to price. Even if the synthesis price falls by two orders of magnitude, amount saved is still substantial and on the order of tens of thousands of dollars.

With respect to the decoder, the time complexity is 𝓞(*K* log *K*) as in Luby Transform. In principle, we could potentially reduce the decoding complexity to 𝓞(*K*) by using Raptor codes[*38*] on top of the Luby Transform. We decided not to pursue this option since Raptor codes are patented and we were concerned that the legal complexities of using this patent could hamper adoption of our method.

### 1.4 DNA Fountain experiments

#### 1.4.1 Computing

All encoding and decoding experiments were done using a MacBook Air with a 2.2 GHz Intel Core i7 and 8Gbyte of memory. The code was tested with Python 2.7.10.

#### 1.4.2 Software, data, and input files

Code, data, and input files are available on:

http://dnafountain.teamerlich.org/

https://github.com/TeamErlich/dna-fountain

Figure S8 presents snapshots of some of the input files in our library.

**Figure S8:**
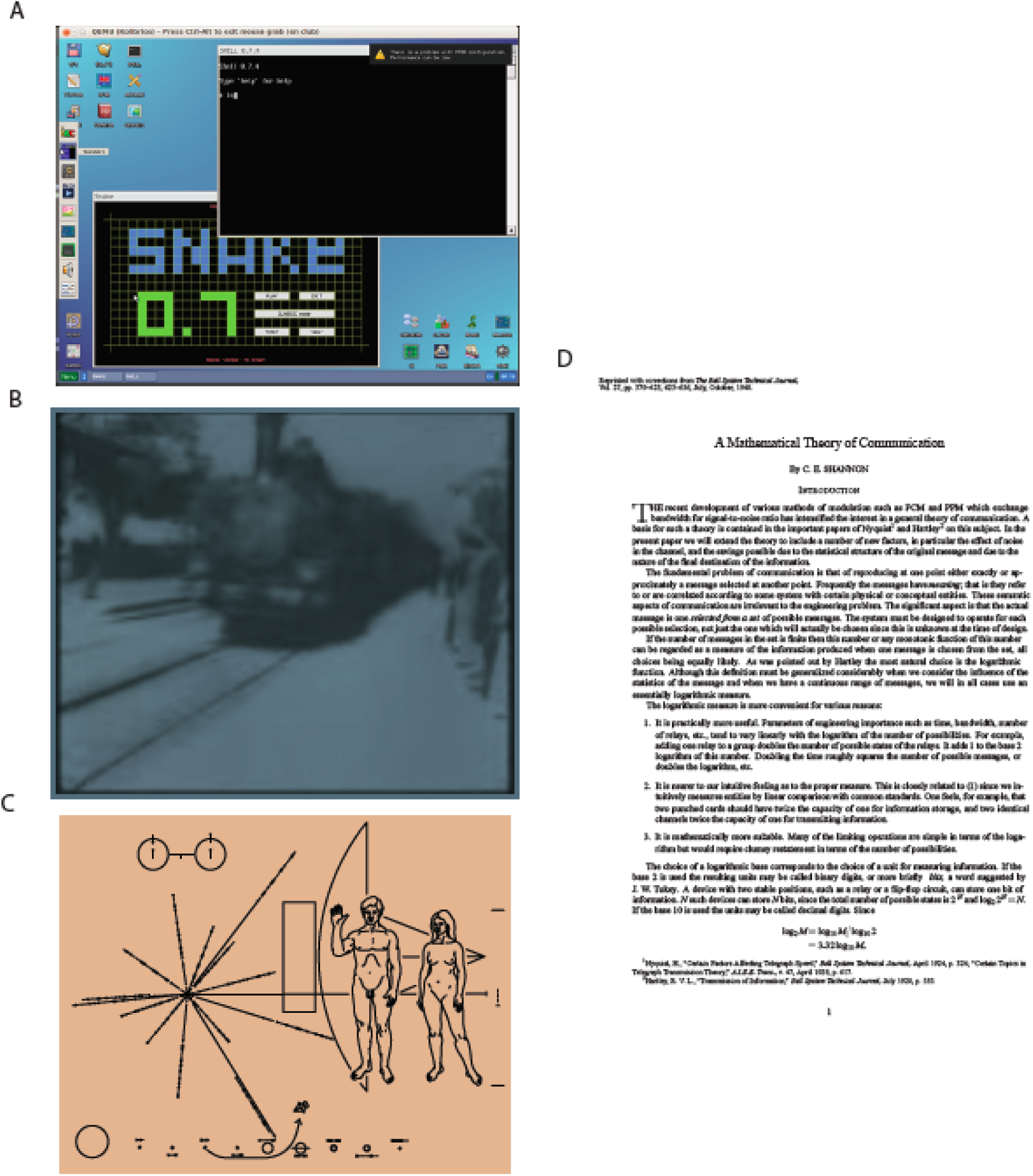
Input files encoded on DNA (A) The Kolibri operating system (B) The Arrival of a Train (C) The Pioneer plaque (D) Shannon’s manuscript on information theory.

#### 1.4.3 Command line steps to encode the data

For reproducibility, we provide the step-by-step commands:

~~~
# Packing input files to a tar and compressing:
tar -b1 -czvvf info_to_code.tar.gz./info_to_code/

#Zero-padding to make the input be a multiple of 512bytes
truncate -s2116608./info_to_code.tar.gz

# Actual encoding of data as DNA:
python encode.py \
–file_in info_to_code.tar.gz \
–size 32 \
-m 3 \
–gc 0.05 \
–rs 2 \
–delta 0.001 \
–c_dist 0.025 \
–out info_to_code.tar.gz.dna \
–stop 72000
–output is a FASTA file named info_to_code.tar.gz.dna

# Adding annealing sites:
cat info_to_code.tar.gz.dna | \
grep -v ’>’ |
awk '{print "GTTCAGAGTTCTACAGTCCGACGATC"$0"TGGAATTCTCGGGTGCCAAGG"}' \
> info_to_code.tar.gz.dna_order

# Sent info_to_code.tar.gz.dna_order to synthesis company
~~~

#### 1.4.4 Molecular procedures

The oligo pool was synthesized by Twist Bioscience. The lyophilized pool consisted of 72,000 oligos of 200nt, which included the 152nt payload flanked by landing sites for sequencing primers:

GTTCAGAGTTCTACAGTCCGACGATC[N152]TGGAATTCTCGGGTGCCAAGG

The pool was resuspended in 20uL TE for a final concentration of 150 ng/ul. PCR was performed using Q5 Hot Start High-Fidelity 2X Master Mix (NEB # M0494) and Illumina small RNA primers RP1 and RPl1 (100ng oligos, 2.5ul of each primer (10µM), 25ul Q5 Master Mix in a 50ul reaction).

PCR Primer (RP1):

5’ AATGATACGGCGACCACCGAGATCTACACGTTCAGAGTTCTACAGTCCGA

PCR Primer, Index 1 (RPI1):

5’ CAAGCAGAAGACGGCATACGAGATCGTGATGTGACTGGAGTTCCTTGGCACCCGAGAATTCCA

Thermocycling conditions were as follows: 30s at 98C; 10 cycles of: 10s at 98C, 30s at 60C, 30s at 72C, followed by a 5 min. extension at 72C. The library was then purified in a 1:1 Agencourt AMPure XP (Beckman Coulter # A63880) bead cleanup and eluted in 20ul water. This library was considered the master pool. Running the library on a BioAnalyzer showed an average size of 263nt (**Figure S9)**, close to the expected 265nt expected by the oligo length (152nt before annealing sites) and the two PCR adapters of length 50nt and 63nt.

**Figure S9:**
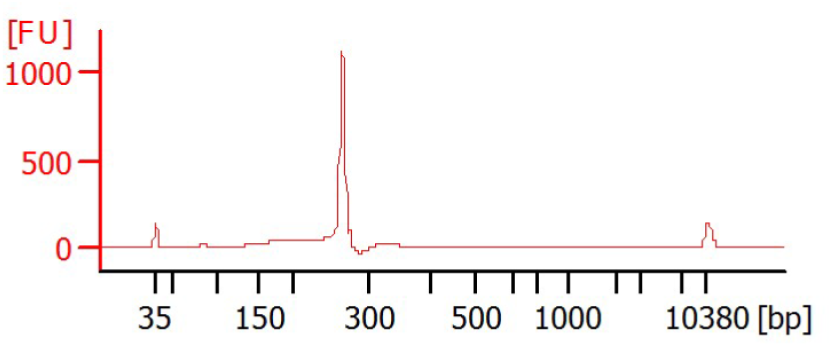
BioAnalyzer results for the oligo pool. The average fragment size was 263nt, concordant with an oligo length of 200nt and the two PCR adapters.

We sequenced the oligo pool using on a single flow cell of the Miseq v3 kit (Illumina # MS-102-3001) at the New York Genome Center using a a 150 pair-end read protocol. Raw .bcl files were downloaded from BaseSpace and analyzed using the commands below.

#### 1.4.5 Decoding the data

~~~
#Converting bcl to fastq using picard (https://github.com/broadinstitute/picard):
for i in {1101̈1119} {2101̈2119}; do
  mkdir ~/Downloads/fountaincode/seq_data3/$i/;
done
for i in {1101̈1119} {2101̈2119}; do
  java -jar ~/Downloads/picard-tools-2.5.0/picard.jar \
  IlluminaBasecallsToFastq \
  BASECALLS_DIR=./raw/19854859/Data/Intensities/BaseCalls/ \
  LANE=1 \
  OUTPUT_PREFIX=./seq_data3/$i/ \
  RUN_BARCODE=19854859 \
  MACHINE_NAME=M00911 \
  READ_STRUCTURE=151T6M151T \
  FIRST_TILE=$i \
  TILE_LIMIT=1 \
  FLOWCELL_BARCODE=AR4JF;
done

#Read stitching using PEAR (Zhang J et al., Bioinformatics, 2014).
This step takes the 150nt reads and places them together to get back the full oligo.
for i in {1101̈1119} {2101̈2119}; do
  pear -f ./$i.1.fastq -r ./$i.2.fastq -o $i.all.fastq;
done
#Retain only fragments with 152nt as the original oligo size:
awk ’(NR%4==2 && length($0)==152){print $0}’ \
*.all.fastq.assembled.fastq > all.fastq.good
#Sort to prioritize highly abundant reads:
sort -S4G all.fastq.good | uniq -c > all.fastq.good.sorted

gsed -r ’s/"\s+//’ all.fastq.good.sorted |\
sort -r -n -k1 -S4G > all.fastq.good.sorted.quantity
#Exclude column 1 that specifies the number of times a read was seen and exclude reads
  with N:
cut -f2 -d’ ’ all.fastq.good.sorted.quantity |\
grep -v ’N’ > all.fastq.good.sorted.seq

#Decoding:
python ~/Downloads/fountaincode/receiver.py \
-f ./seq_data3/all.fastq.good.sorted.seq \
–header_size 4 \
–rs 2 \
–delta 0.001 \
–c_dist 0.025 \
-n 67088 \
-m 3 \
–gc 0.05 \
–max_hamming 0 \
–out decoder.out.bin

#Verification:
md5 decoder.out.bin
#output is 8651e90d3a013178b816b63fdbb94b9b
md5 info_to_code.tar.gz
#output is 8651e90d3a013178b816b63fdbb94b9b

#YES!
~~~

#### 1.4.6 Creating a deep copy

For the nine consecutive PCR reactions, we started with the master pool as described above. Then, we performed 9 subsequent rounds of PCR using the same conditions as above. The first round used 10ng of oligo input from the master pool into 25ul PCR reactions. Then, 1ul from each PCR reaction was input into each subsequent round of PCR for a total of 9 reactions without cleanup. The final round was purified in a 1:1 Agencourt AMPure XP bead cleanup and eluted in 20ul deionized water. The final library was run under the same conditions as described.

This procedure can theoretically create 300 × 25^9^ × 2 = 2.28 × 10^15^ copies of the file (see **Figure S11)**. The decoding procedure for this library is identical to the one presented for the master copy. Fitting the negative bionomial distribution was done in R using the fitdistr command.

**Figure S10:**
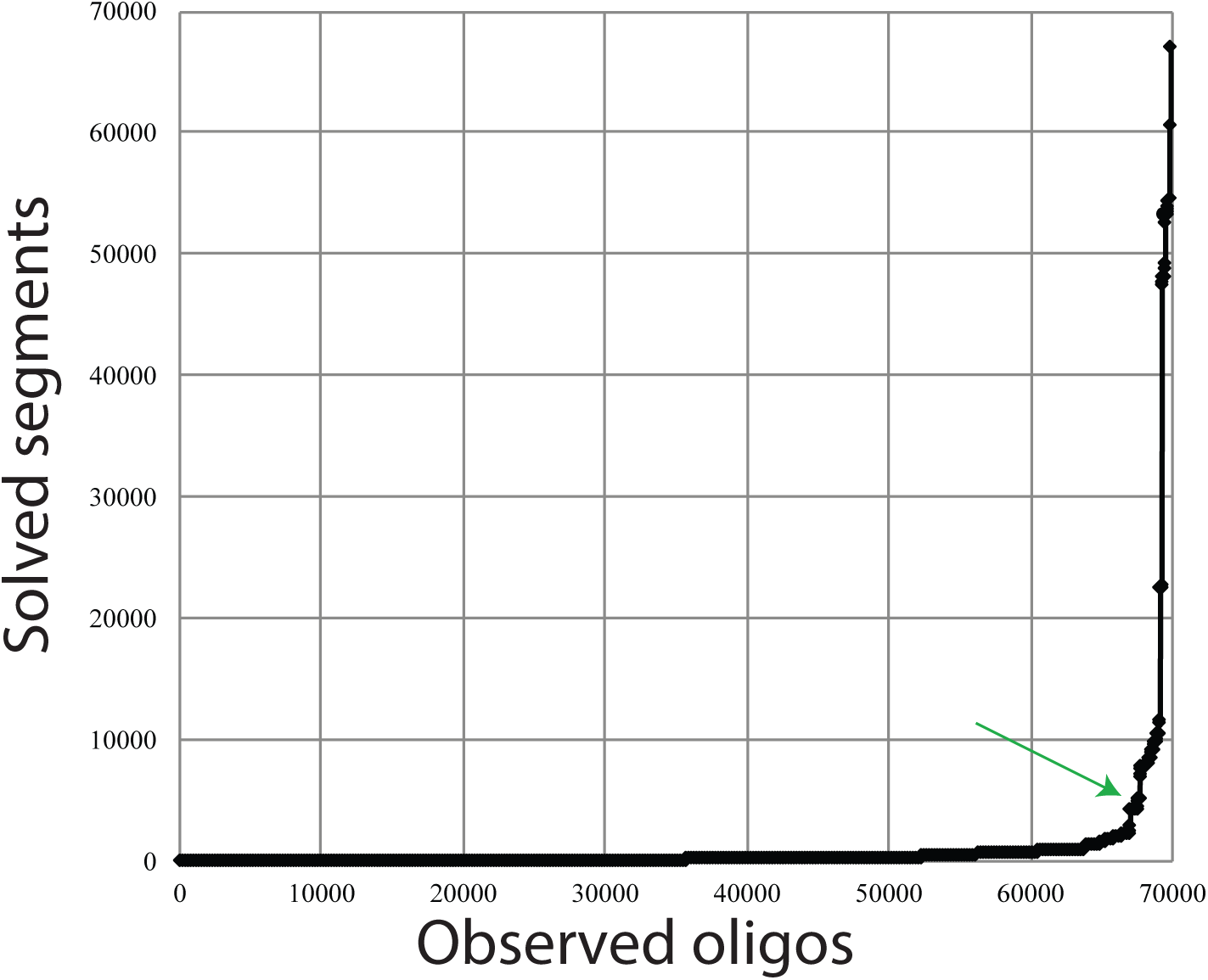
The number of solved segments as a function of observed oligos. The figure demonstrates the avalanche process. At first, the observed oligos cannot determine the data in the input segments. When the number of observed oligos is about the same as the number of input segments (green arrow), sufficient information has accumulated to infer the data in the input segments and an avalanche of inference starts with each new oligo. Notice that the decoder needed only to observe 69800 oligos out of the 72,000 synthesized oligos to decode the entire file, illustrating the robustness against dropouts.

**Figure S11:**
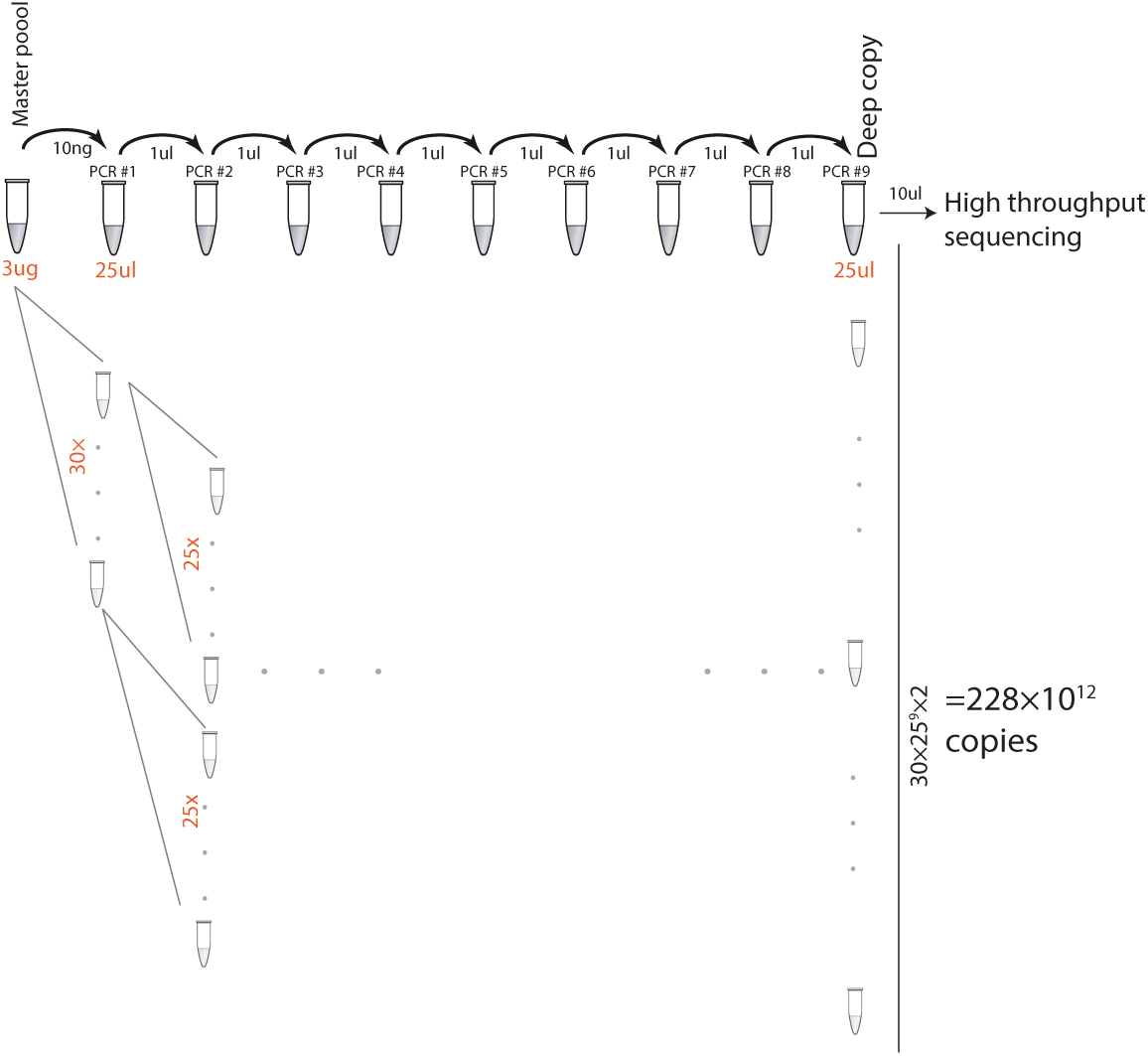
Exponential amplification process. Each PCR reaction generates enough volume for multiple subsequent reactions. The first PCR reaction used only 10ng from the master pool of 3ug of DNA. This implies that 300 similar reactions can be conducted using the master pool. Each subsequent PCR used 1ul out of 25ul generated by the previous reaction. Therefore, an exponential process could amplify the number of copies by 25 times in each reaction (gray tubes). Eventually, we consumed about half of the final reaction for sequencing and QC, meaning that the final reaction can be sequenced twice. In total, the nine step amplification process has the potential to create 300 × 25^9^ × 2 = 2.28 quadrillion copies. Our experiment reflects one end-to-end arm of this copy (black tubes).

#### 1.4.7 Determining the lowest number of molecules

The Church et al. prediction regarding theoretical density of DNA storage was derived from their main text (5.5 petabit/mm3) and using their estimated mass density of DNA of 1mg/1mm3.

To generate a serial dilution of the DNA Fountain library, we started with the original library generated by Twist. The concentration of this library was determined by Nanodrop single-strand DNA measure and was found to be 150ng/ul. To generate condition 1, we obtained 2ul of DNA from this library into 28 of TE buffer. Measuring condition 1 using Nanodrop confirmed that the concentration was 10ng/ul. Next, we diluted the library by a factor of ×10 by taking 4ul of the previous condition into 36ul of TE and repeating this procedure 6 more times to generate libraries in concentrations of 10ng, 1ng, 100pg, 10pg, 1pg, and 100fg, and 10fg. We then took 1ul of each library to a PCR reaction of 50ul and run the reaction for 40cycles. Each library was barcoded with a distinct 6nt barcode to allow multiplexing. We then loaded the libraries using the Illumina standard protocol and sequenced the libraries using two lanes of the Rapid Sequencing kit to 150 pair-end reads.

The number of molecules per oligo for each dilution was determined according to: *w*_*i*_[g]/(200[nt] ×325[Dalton/nt] ×1.67 × 10^−24^[g/Dalton] ×72000[oligos]), where *w*_*i*_ is the total weight of DNA in the library (e.g. 10ng, 1ng, …).

The decoding command was:

~~~
#generating fountain-4.all.fastq.good.sorted.quantity is done using the pipeline
described above.
cat fountain-4.all.fastq.good.sorted.quantity | python receiver.py -f - –header_size 4
  –rs 2 –delta 0.001 –c_dist 0.025 -n 67088 -m 3 –gc 0.05 –max_hamming 0 –out
  decoder.out.bin.tar.gz –kmer 20
~~~

To test the unrealistic decoder, we scanned all sequence reads that passed assembly regardless of their length. Next, if we found a unique match of at least 15nt to one of the 72000 oligos, we replaced the read with the original oligo. Finally, we counted the number of unique oligos. In all cases, we could get enough unique oligos for decoding.

